# Lysinoalanine crosslinking is a conserved post-translational modification in the spirochete flagellar hook

**DOI:** 10.1101/2023.06.13.544825

**Authors:** Michael J. Lynch, Maithili Deshpande, Kurni Kyrniyati, Kai Zhang, Milinda James, Michael Miller, Sheng Zhang, Felipe J. Passalia, Elsio A. Wunder, Nyles W. Charon, Chunhao Li, Brian R. Crane

## Abstract

Spirochete bacteria cause Lyme disease, leptospirosis, syphilis and several other human illnesses. Unlike other bacteria, spirochete flagella are enclosed within the periplasmic space where the filaments distort and push the cell body by action of the flagellar motors. We previously demonstrated that the oral pathogen *Treponema denticola* (Td) catalyzes the formation of covalent lysinoalanine (Lal) crosslinks between conserved cysteine and lysine residues of the FlgE protein that composes the flagellar hook. Although not necessary for hook assembly, Lal is required for motility of Td, presumably due to the stabilizing effect of the crosslink. Herein, we extend these findings to other, representative spirochete species across the phylum. We confirm the presence of Lal crosslinked peptides in recombinant and *in vivo*-derived samples from *Treponema* spp., *Borreliella* spp., *Brachyspira* spp., and *Leptospira* spp.. Like with Td, a mutant strain of the Lyme disease pathogen *Borreliella burgdorferi* unable to form the crosslink has impaired motility. FlgE from *Leptospira* spp. does not conserve the Lal-forming cysteine residue which is instead substituted by serine. Nevertheless, *Leptospira interrogans* also forms Lal, with several different Lal isoforms being detected between Ser-179 and Lys-145, Lys-148, and Lys-166, thereby highlighting species or order-specific differences within the phylum. Our data reveals that the Lal crosslink is a conserved and necessary post-translational modification across the spirochete phylum and may thus represent an effective target for spirochete-specific antimicrobials.

**Significance Statement:** The phylum Spirochaetota contains bacterial pathogens responsible for a variety of diseases, including Lyme disease, syphilis, periodontal disease, and leptospirosis. Motility of these pathogens is a major virulence factor that contributes to infectivity and host colonization. The oral pathogen *Treponema denticola* produces a post-translational modification (PTM) in the form of a lysinoalanine (Lal) crosslink between neighboring subunits of the flagellar hook protein FlgE. Herein, we demonstrate that representative spirochetes species across the phylum all form Lal in their flagellar hooks. *T. denticola* and *B. burgdorferi* cells incapable of forming the crosslink are non-motile, thereby establishing the general role of the Lal PTM in the unusual type of flagellar motility evolved by spirochetes.

## Introduction

Pathogenic spirochetes cause a myriad of human and animal diseases, including Lyme disease, periodontal disease, syphilis, pinta, yaws, endemic syphilis, leptospirosis, swine dysentery, and bovine digital dermatitis.^1–10^ Currently, the phylum Spirochaetota is summarized as a single class, Spirochaetia, that is subdivided into four orders: Spirochaetales, Brevinematales, Brachyspirales, and Leptospirales (Figure 1A).^11,12^ Although this classification scheme is relatively mature, as more spirochete species are identified, cultured, and their genomes sequenced, taxonomic organization continues to be updated for this complex collection of bacterial species.^12–14^

**Figure 1.**
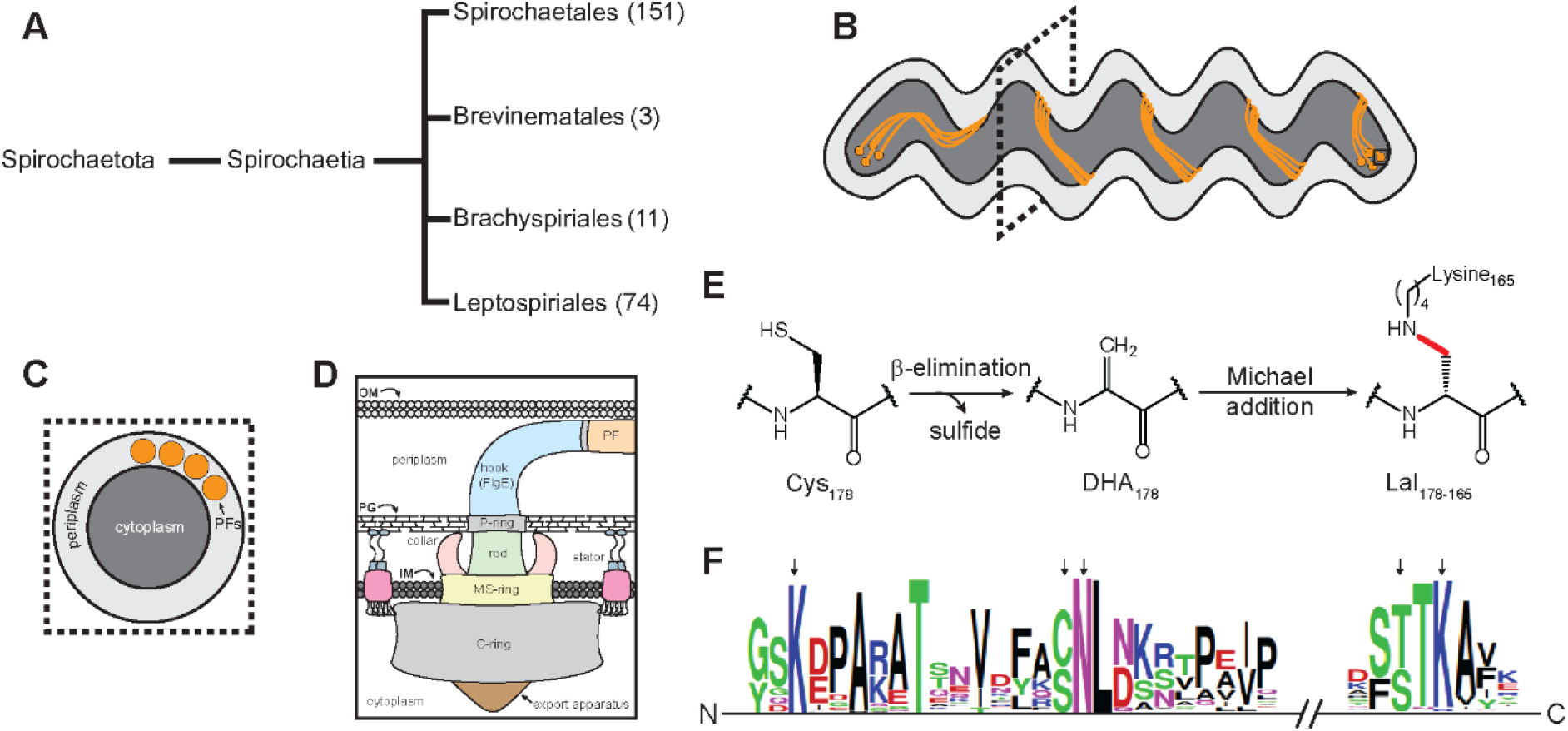
Lysinoalanine (Lal) crosslinking in the spirochete flagellar hook protein FlgE. **(A)** Simplified taxonomic organization of the Spirochaetota phylum. The phylum is organized into a single class, which is further divided into four distinct orders. The approximate number of species present in each order is included for comparison (excluding environmental, unclassified, and uncultured species). (**B)** Cartoon depiction of a spirochete cell with multiple subpolar membrane-embedded flagellar motors (solid black square) linked by a continuous ribbon of periplasmic filaments (PFs). (**C)** Cross-section of the spirochete cell body as the position indicated by the dotted square in B. The four PFs wrap around the cell body. (**D)** Overview of the sub-structure of the bacterial flagellar motor (OM -outer membrane, PG – peptidoglycan, IM – inner membrane). (**E)** Biochemical mechanism of Lal crosslink formation in the flagellar hook protein FlgE of *T. denticola* (Td). Residue numbering is based on the *T. denticola* FlgE (Lal crosslink indicated by red line). (**F)** Multiple sequence alignment of the 241 FlgE sequences collected and filtered using Annotree^60^. Black arrows (left to right, Td numbering) indicate the position of Lys-165, Cys-178, Asn-179, Thr-334, and Lys-336 residues. Sequence graphics created using WebLogo.^62^

Similar to other bacteria, spirochetes rely on chemotaxis to detect and migrate towards attractants (e.g., sugars, amino acids, rabbit serum, long-chain fatty acids, and hemoglobin) and away from repellants (e.g., ethanol, butanol).^15–21^ Motility is a widely recognized virulence factor for pathogenic spirochetes.^3,7,22– 24^ Similar to other flagellated bacteria, spirochete motility is powered by membrane-embedded rotary motors connected to helical flagellar filaments (Figure 1B-D).^25^ However, unlike the majority of flagellated bacteria which have external flagella like *Escherichia coli* and *Salmonella* spp., spirochetes enclose their flagellar filaments in the periplasm.^7^ These periplasmic filaments (PFs) wrap around the cell body, and in some species, form a continuous ribbon of overlapping PFs from pole to pole (Figure 1B-C). To push the cell forward, PF rotation in the periplasm deforms the cell body, inducing undulations that travel as rolling waves along the cell body and producing thrust. Interestingly, in some species, PFs also serve a cytoskeletal role, warping the cell body to yield their distinct flat-wave or corkscrew morphology.^1^ PFs are connected to the motor via the hook (Figure 1D). The hook is a highly curved, helically tubular structure comprised of ∼130 subunits of a single protein FlgE.^26^ Within the motor, the hook serves to transmit motor torque to the PFs. As a result, the hook must be flexible enough to rotate within the periplasm yet strong enough to transfer torque to the filament-wrapped cell body and avoid buckling.^27^ FlgE monomers organize into 11 protofilaments that further assemble into a right-handed helix.^28^ To provide the flexibility required for rotation, extensive networks of protein-protein contacts between neighboring FlgE subunits strengthen the hook, but also allow change depending on the rotational state of the protofilament.

Although the structure of the bacterial flagella is generally well conserved across the bacterial kingdom, species-specific differences are common.^25^ In spirochetes, these include such examples as the addition a motor component known as the P-collar^29^, glycosylation of the PFs^30^, the use of multiple flagellin and sheath proteins in the PFs^31–34^, and a lysinoalanine (Lal) post-translational modification (PTM) found in FlgE.^35–37^ FlgE from *Treponema denticola* (Td) and *Borreliella burgdorferi* (Bb) self-catalyze the formation of a Lal crosslink between conserved cysteine and lysine residues (Figure 1D) harbored on adjacent FlgE subunits within the hook.^35,36^ Unlike other forms of Lal found in nature, Lal formation in Td FlgE is autocatalytic and will occur in the absence of other enzymes, cofactors, or regulatory proteins.^38^ FlgE Lal crosslinking follows at least three distinct biochemical steps (Figure 1D). First, TdFlgE oligomerizes via protein-protein interactions between long, highly conserved N- and C-terminal α-helical D0 domains.^35^ β-elimination of the catalytic cysteine yields the reactive dehydroalaine (DHA) intermediate, producing hydrogen sulfide as a byproduct.^35,39^ Lastly, DHA reacts with the ε-NH2 group of lysine-165 in a Michael addition reaction to yield the mature Lal crosslink (red line in Figure 1D). To date, we confirmed the presence of Lal crosslinked peptides in recombinantly-derived *T. denticola* FlgE and polyhook PFs from an engineered *B. burgdorferi* strain lacking the hook length regulatory protein FliK.^36^ Mutagenesis of catalytic residues C178, K165, and structural residue N179 inhibited Lal formation in *T. denticola* cells *in vivo* and rendered the cells non-motile.^36^

Although *T. denticola and B. burgdorferi* only represent one of the four orders that comprise the phylum Spirochaetota, multiple-sequence alignment data from spirochetes across the phylum suggest that Lal crosslinking is a conserved PTM (Figure 1E). Herein, we test this hypothesis directly, detecting Lal crosslinking in *in vitro-* and *in vivo-*derived samples from *Treponema* spp., *Borreliella* spp., *Brachyspira* spp., and *Leptospira* spp.. These species span genera, families, and represent three out of the four orders in the phylum. Interestingly, our findings also reveal that *Leptospira interrogans* catalyzes multiple different Lal crosslinks within the hook, a polymorphism likely conserved by other *Leptospira* spp. based on sequence conservation. Finally, swimming speed analysis of *B. burgdoferi* cells harboring the C178A mutation in FlgE demonstrates the requirement of the Lal crosslinking for motility in this species as well as *T. denticola*. Overall, these findings suggest that Lal crosslinking is a conserved PTM of the spirochete flagellar hook that is required for proper hook function and optimal motility.

## Results

### Treponema denticola and Borreliella burgdorferi as model systems for identifying Lal crosslinking

For this study, we initially aimed to confirm the presence of Lal-crosslinked FlgE peptides in unmodified spirochete strains. However, due to the challenging culturing conditions of many spirochete species, this was not feasible for certain spirochetes. As a result of these constraints, as well as the ability of FlgE proteins to self-catalyze the crosslink, we also explored recombinant samples when and/or if obtaining a wild-type (WT) PF sample was not possible (Fig. 2A). *In vivo* samples (Fig. 2A. top) originated from either WT PFs, or from polyhook (PH) PFs. PH PFs were purified from engineered spirochete cells harboring a *fliK* gene deletion (*fliK*Δ) modification that removes the ability of cells to regulate their look lengths. As a result, fliKΔ cells express elongated hooks that are several times longer than normal flagellar hooks.^40^ This approach was employed to increase the FlgE concentration compared to WT PF samples, thus increasing the probability of successfully identifying Lal-crosslinked peptides via mass spectrometry (MS). If the WT or fliKΔ spirochete cells could not be cultured, the *flgE* gene was overexpressed in *E. coli* cells (Figure 2, bottom). The proteins were then purified and incubated in crosslinking buffer to drive *in vitro* Lal crosslink formation.^35,36^ Regardless of the source of FlgE protein, all sample types (WT, PH, and recombinant FlgE) were treated identically following *in vivo* PF isolation or *in vitro* Lal crosslinking assays. Previous studies have shown that endogenous FlgE proteins from *T. denticola* (Td) and *B. burgdorferi* (Bb) form high-molecular weight complexes (HMWCs) consisting of Lal crosslinked FlgE monomers when visualized vis SDS-PAGE or western blot.^36,37^ These HMWCs run at the top of SDS-PAGE gels, regardless of sample treatment prior to SDS-PAGE.^36,37^ Therefore, recombinant or *in vivo*-derived FlgE samples were denatured, electrophoresed and only the top region of the gel was extracted and submitted for HPLC-MS analysis (dotted box in Figure 2). To detect Lal-containing peptides by MS, we employed a similar procedure described previously for recombinant TdFlgE and PH BbFlgE.^36^To address whether Lal is a conserved PTM present in species throughout the phylum Spirochaetota, we first examined Td and Bb FlgE (Figure 3A-D) to demonstrate that recombinant FlgE crosslinking recapitulates the same Lal product as found in PH and/or WT samples. These two species belong to the order spirochaetales but reside in different taxonomic families (Treponemataceae versus Borreliella, respectively). The Lal-containing typtic-peptides were confirmed in WT Td derived PFs (Figure 3A and Fig. S1). The extracted-ion chromatogram (XIC) and the electron-transfer dissociation (ETD) MS/MS fragmentation data agree with the expected mass and c/z fragmentation pattern of the parent Lal-peptide. Taken together with our previous findings, we show that Lal-crosslinked FlgE peptides are present in all three sample types tested (WT PFs, PH PFs, and recombinant TdFlgE). A similar approach with Bb FlgE samples produced similar results, with XIC and MS/MS data confirming the presence of Lal crosslinked peptides in both *in vitro-* and *in vivo-*derived Bb FlgE samples (Figure 3B-D and SI Figure 2-4). Overall, in the test cases of Td and Bb, crosslinked peptides derived from three different FlgE sources related to the same strain all contain the same Lal species.

**Figure 2.**
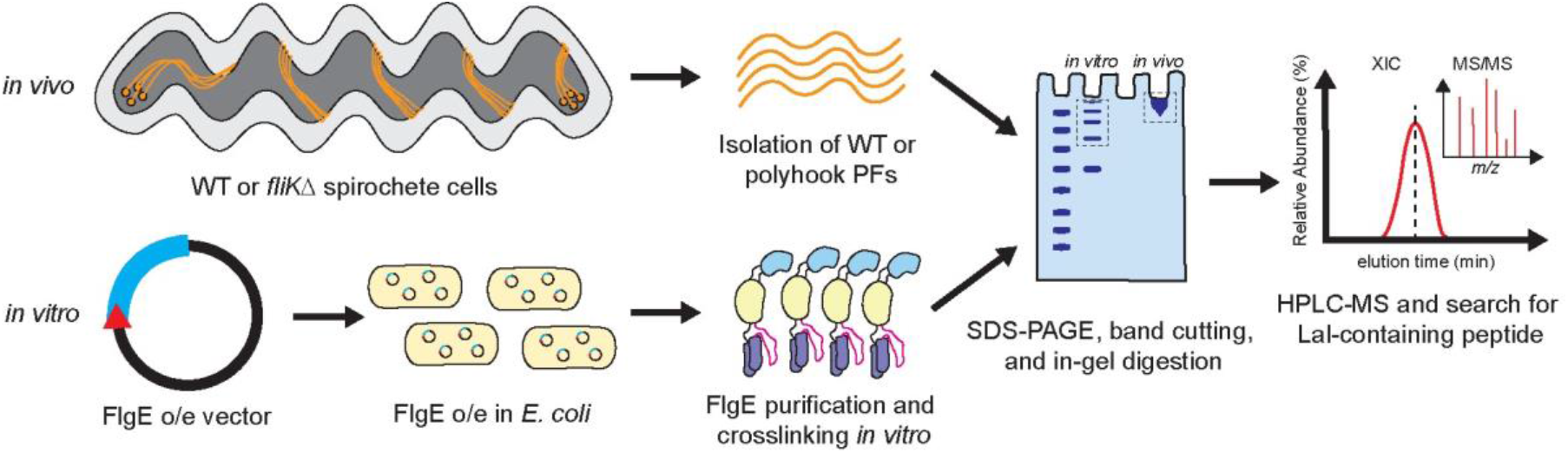
Overview of sample types used in this study. (top) *In vivo* FlgE samples were derived from either WT or *fliK* knock-out (*fliK*Δ) cell lines. (bottom) *In vitro* samples were generated via recombinantly expressing spirochete FlgE proteins in *E. coli* cells and performing Lal cross-linking assays prior to mass-spec analysis. All sample types were run on SDS-PAGE, and the appropriate high molecular weight bands (dashed boxes) excised and submitted to MS for Lal detection.

**Figure 3.**
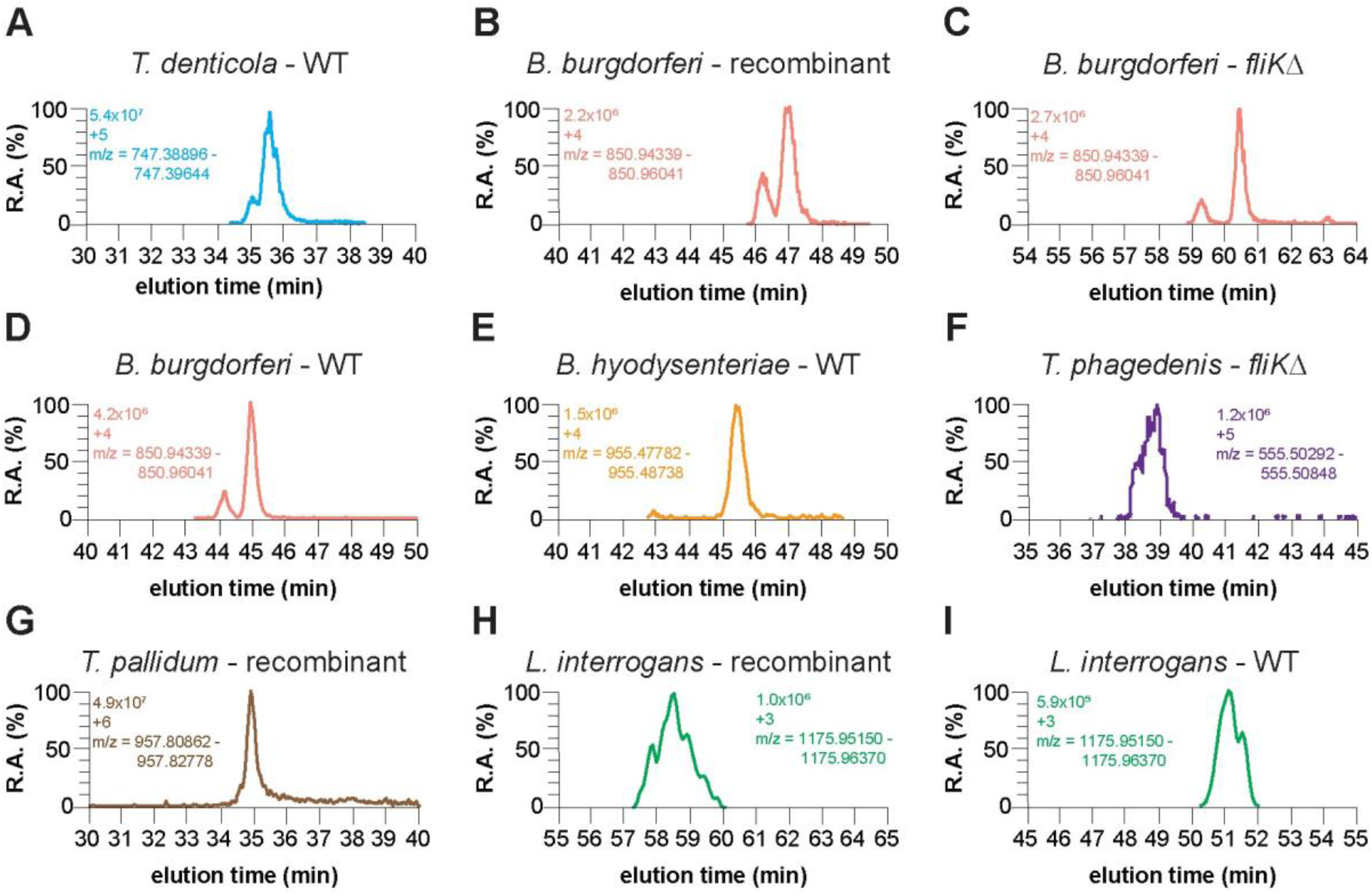
Detection of Lal crosslinked FlgE in recombinant, *fliK*Δ, and wild-type cell culture samples. Extracted-ion chromatogram (XIC) of **(A)** *T. denticola* WT PFs. **(B)** *B. burgdorferi* recombinant FlgE, **(C)** *B. burgdorferi fliK*Δ, **(D)** *B. burgdorferi* WT PFs, **(E)** *B. hyodysenteriae* WT PFs, **(F)** *T. phagedenis flik*Δ, **(G)** *T*.*pallidum* recombinant FlgE, **(H)** *L. interrogans* recombinant FlgE, and **(I)** *L. interrogans* WT PFs Lal crosslinked FlgE peptides. For each XIC, the intensity levels, peptide charged state, and m/z range are shown.

### Detection of FlgE Lal crosslinks in diverse pathogenic spirochetes

We next sought to confirm the presence of Lal crosslinked FlgE peptides in other, more distantly related pathogenic spirochetes. We first examined the spirochete pathogen associated with swine dysentery, *Brachyspira hyodysenteriae* (Bh, formerly known as *Treponema* or *Serpulina*).^41^ Bh belongs to the order Brachyspirales (Figure 1A). Bh FlgE conserves lysine (149) and cysteine (162) residues analogous to those that form Lal (Lys165 and Cys178, respectively) in Td and Bb. Indeed, following cultivation of Bh and isolation of PFs, the XIC yielded peptides eluting at 61.08 minutes in the [M+4H]^4+^ and [M+5H]^5+^ charged state matched the expected m/z of the Lal crosslinked Bh FlgE tryptic peptide (Figure 3E and SI Figure 5). The ETD MS/MS fragmentation of this peptide produced y and b ions with m/z values consistent with the expected Lal crosslinked peptide (SI Figure 5).

In addition to the periodontal disease and gingivitis-associated pathogen Td, the genus Treponema has other notable pathogens that pose risks to humans and other mammals. These include the two *Treponema* species we examine here: *Treponema phagedenis* strain Kazan 5 (Tph) and *Treponema pallidum* subspecies *pallidum* strain Nichols (Tpa). *Treponema phagedenis-*like species are associated with bovine digital dermatitis.^8^ Inspection of the Tph FlgE primary sequence suggests that it also catalyzes Lal crosslinks (Figure 1E and SI Figure 6A). Isolation and MS analysis of PH PFs from Tph fliKΔ cells yielded peptides in the [M+4H]^4+^ and [M+5H]^5+^ charged states that eluted between 37.6 -37.7 minutes (Figure 3F and SI Figure 6). The ETD fragmentation pattern of the [M+5H]^5+^ peptide produced c and z ions with m/z values consistent with the parent Lal crosslinked peptide, confirming the presence of Lal crosslinking in TphFlgE (SI Figure 6).

Tpa is one of four *Treponema pallidum* subspecies that are associated with human disease and is the pathogen responsible for venereal syphilis. The other three related strains are the causative agents of other human diseases like yaws (*Treponema pallidum* subsp *pertenue*), endemic syphilis (*Treponema pallidum* subsp *endemicum*), and pinta (*Treponema carateum*).^42^ Although difficult to culture *in vitro*,^42–44^ virulent Tpa cells were cultured *in vitro* and isolated from rabbit testes in collaboration with Steve Norris and Diane Edmondson, University of Texas, McGovern Medical School, Houston using their recently developed cell culture methodology.^45–47^ Unfortunately, the concentration of Tpa FlgE in the final WT PF samples was too low to confidently detect Tpa FlgE and any Lal crosslinked tryptic peptides. Given our previous data with Td and Bb FlgE, we produced Tpa FlgE recombinantly in *E. coli* and tested it for Lal crosslinking. *In vitro* crosslinking of Tpa FlgE yielded multimer high molecular weight bands that formed in a time-dependent manner, albeit at a lower extent compared to similar concentrations of Td FlgE. Excision and in-gel digestion of the multimer bands yielded a peak on the MS XIC with an m/z range that corresponded to a Tpa FlgE Lal crosslinked-peptide in a [M+6H]^6+^ charged state (Figure 3G). The Tpa FlgE Lal peptide was also detected in two other charged states ([M+5H]^5+^ and [M+7H]^7+^) that all eluted at ∼34.9 minutes (SI Figure 7B). ETD MS/MS spectra of the [M+6H]^6+^ peptide produced c and z ions that were consistent with the parent Lal crosslinked peptide (Figure 2F and SI Figure 7A and 7C).

The order Leptospirales contains FlgE proteins that are the most diverse within the spirochetes with respect to Lal crosslinking. We examined *Leptospira interrogans* serovar Copenhageni strain Fiocruz L1-130. *L. interrogans* belongs to the order Leptospirales (Figure 1A) and is one of over 41 species and >300 serovars of pathogenic Leptospira responsible for the human and non-human disease known as leptospirosis.^4,48–51^ Although several serogroups are associated with leptospirosis, serovar Copenhageni is recognized as one of the most virulent pathogenic species of Leptospira.^4^ Examination of the *L. interrogans* (Li) strain Fiocruz L1-130 FlgE primary sequence reveals that in addition to the conserved lysine residue, there is a notable difference in the Lal crosslink parent residues compared to the other spirochete species investigated previously. The conserved cysteine residue in Li FlgE has been replaced with a serine residue (SI Figure 8A and Figure 4A-B). Serine has been shown to participate in Lal formation in food processing.^52^ In addition, we have shown that Td FlgE C178S mutant can form Lal crosslinked HMWCs on SDS-PAGE gels in *in vitro* Lal crosslinking assays and that Lal crosslinked peptides are present in these HMWCs via MS.^35^ Thus, we predicted that Li FlgE can also undergo Lal crosslinking. To test this possibility, we analyzed recombinant Li FlgE Lal-crosslinked HMWCs and WT PFs purified from Li cells (Figure 3H-I). XICs of both *in vitro* and *in vivo-*derived Li FlgE samples showed peaks corresponding to Li FlgE Lal-crosslinked peptides in the [M+3H]^3+^ charges state (Figure 3H-I, SI Figure 8 and 9) and the [M+4H]^4+^ and [M+5H]^5+^ state (SI Figure 8B and 9B). ETD fragmentation produced c and z ions that agreed well with the expected c and z ions of the AspN-digested Lal crosslinked peptide, therefore confirming the presence of Lal crosslinking in Li FlgE both *in vitro* and *in vivo* (SI Figure 8C and 9C).

**Figure 4.**
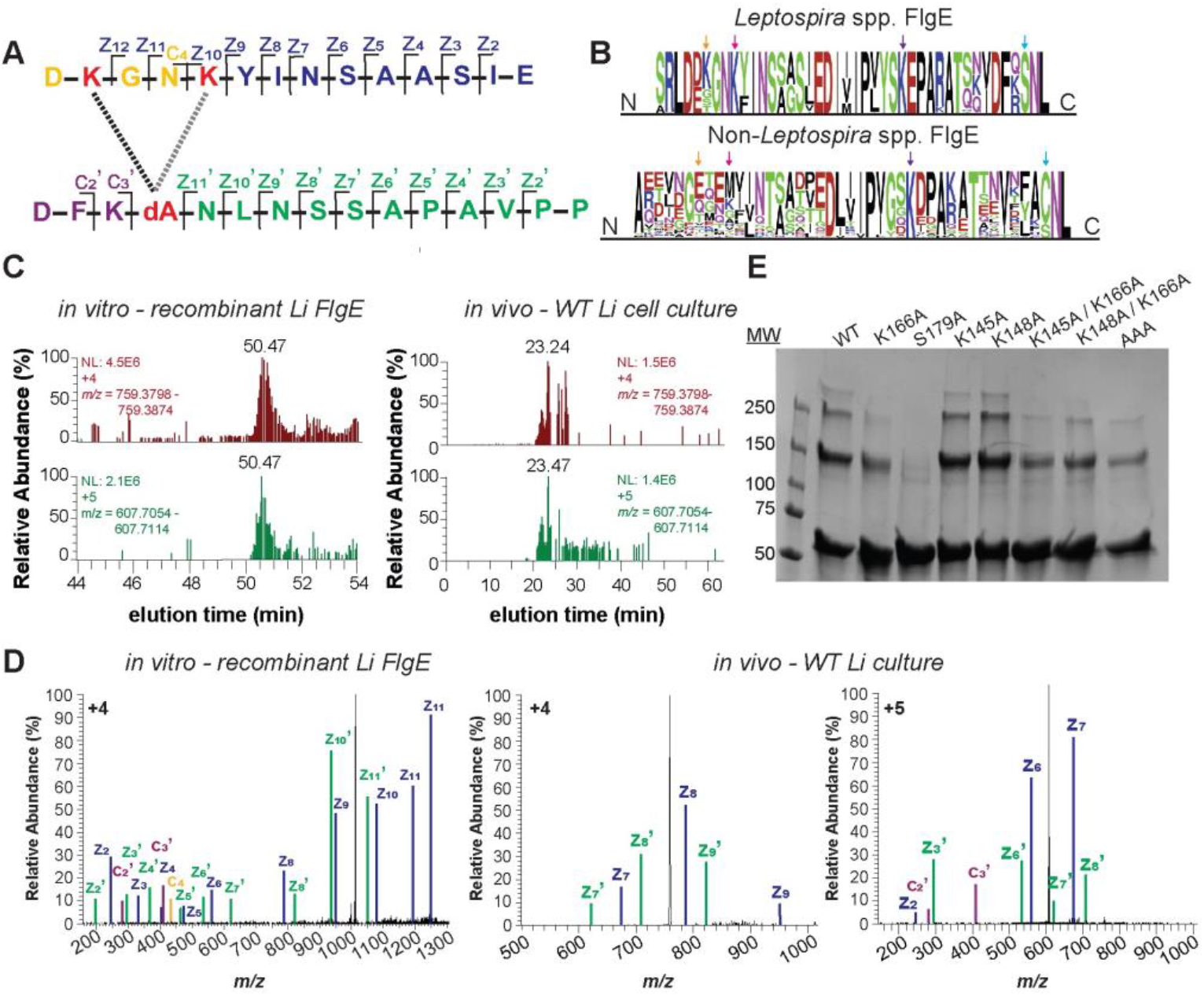
*L. interrogans* FlgE catalyzes multiple Lal crosslinkages. **(A)** Lal crosslinked Li FlgE peptide following AspN digestion. Lys-145, Lys-148, and DHA (dA)-179 are colored red. The two Lal crosslink isoforms are represented by a black (S179-K145) and gray (S179-K148) dotted line. C and Z ions are indicated and color-coded according to the peaks identified in (D). **(B)** Multiple sequence alignment and sequence conservation of Lys-145 (orange arrow, Li numbering) and Lys-148 (pink arrow) in *Leptospira* spp. FlgE (top) versus non-*Leptospira* spp. FlgE (bottom). Purple and blue arrows mark the position of Lys-166 and Ser-179, respectively. **(C)** XICs of recombinant (left) and WT PFs (right) Lal crosslinked Li FlgE peptide. **(D)** MS/MS ETD fragmentation of Lal-crosslinked peptide parent ion with c and z ions annotated and labeled according to (A). MS/MS spectra of *in vitro* recombinant (left, [M+4H]^4+^) and *in vivo* WT cell culture (center and right, [M+4H]^4+^ and [M+5H]^5+^, respectively) Li FlgE Lal-crosslinked peptide. Individual c and z ions were amplified 5-200x. **(E)** *In vitro* SDS-PAGE Lal crosslinking assays of Li FlgE lysine mutants compared to WT.

### L. interrogans FlgE produces several different Lal crosslinks

In addition to the canonical Lal crosslink between K166 and S179 (Figure 3H-I, SI Figure 8-9) in Li FlgE, Lal also forms between S179 and either K145 and/or K148 (Figure 4). These Lal isoforms were detected in both recombinantly expressed *in vitro*-derived Li FlgE and *in vivo* WT Li PFs (Figure 4C, SI Figure 8-9). Due to limited resolution of the MS and the proximity of K145 and K148 in sequence, it was unclear which residue was participating in the alternative crosslink. Sequence conservation of these residues in the order Leptospirales compared to species in the orders Brachyspirales, Brevinematales, and Spirochaetales (Figure 4B) shows that K148 is conserved 100% of the time (67/67 sequences) compared to K145 being conserved 61% of the time (41/67 sequences), thereby suggesting that the Lal isoform more likely involves K148 compared to K145. A similar analysis with non-*Leptospira* spp. reveals that these two residues are highly variable, suggesting that Lal isoforms involving these positions are not common in other orders (Figure 4B). To further verify these findings, we generated alanine substitutions for K145, K148, and K166 and measured the Lal crosslinking ability of these mutants compared to WT LiFlgE in the Lal crosslinking assay.^35^ Compared to WT Li FlgE, the K166A substitution substantially reduced crosslinking, whereas K145A and K148A moderately reduced the yield of HMWCs (Figure 4E). Surprisingly, the K145A/K166A and K148A/K166A double Ala substitutions still formed crosslinked dimers and trimers, which suggested that either K145 or K148 could generate Lal isoforms. Unexpectedly, the triple substitution K145A/K148A/K166A Li FlgE also produced some HMWCs, indicating that yet another lysine residue may have participated in the *in vitro* reaction with S179. Searching the WT PF MS dataset for crosslinks between S179 and other lysine residues did not yield any candidates other than K166, K145 and/or K148 (SI Figure 10A). Furthermore, structural modeling of LiFlgE oligomers using cryo-EM density of *Salmonella* flagellar hooks showed no other lysine residues within reactive distance to S179 (SI Figure 10B).

### Lal crosslinking is required for *Borreliella burgdorferi* motility

Previously, we showed that Lal crosslinking of FlgE is required for motility of the oral pathogen Td. To expand on this finding, we investigated whether Lal crosslinking was required for the motility of another genetically tractable pathogenic spirochete such as Bb. We genetically modified low passage, virulent *B. burgdorferi* strain B31 A3-68 cells to remove the *flgE* gene (*flgE*Δ) or expressed a C178A mutant form of the FlgE protein. In agreement with previous findings from Td, WT and C178A FlgE Bb cells had a flat wave morphology, whereas *flgE*Δ cells were rod-shaped due to the lack of PFs (Figure 5A). The wild-type morphology of Bb C178A FlgE mutant cells confirms the presence of correctly assembled hooks and PFs since PFs serve a cytoskeletal role in Td and Bb. This observation suggests that hook and PF assembly does not depend on the presence of Lal crosslinking between adjacent FlgE subunits. Western blot analysis of whole-cell lysate of each strain confirmed the presence of FlgE HMWCs in WT Bb cells and the absence of FlgE in the *flgE*Δ mutant cells (Figure 5B). Additionally, the Western blot of the C178A FlgE strain showed only a FlgE band at the MW of the single subunit (Figure 5B). Swimming speed analysis of WT (12 ± 2 μm/s, n=24) and C178A (0.3 ± 2 μm/s, n=49) cells reveal that despite having a normal morphology and correctly assembled PFs, the lack of Lal crosslinks between adjacent FlgE monomers in the flagellar hook of Bb C178A cells renders the cells non-motile.

**Figure 5.**
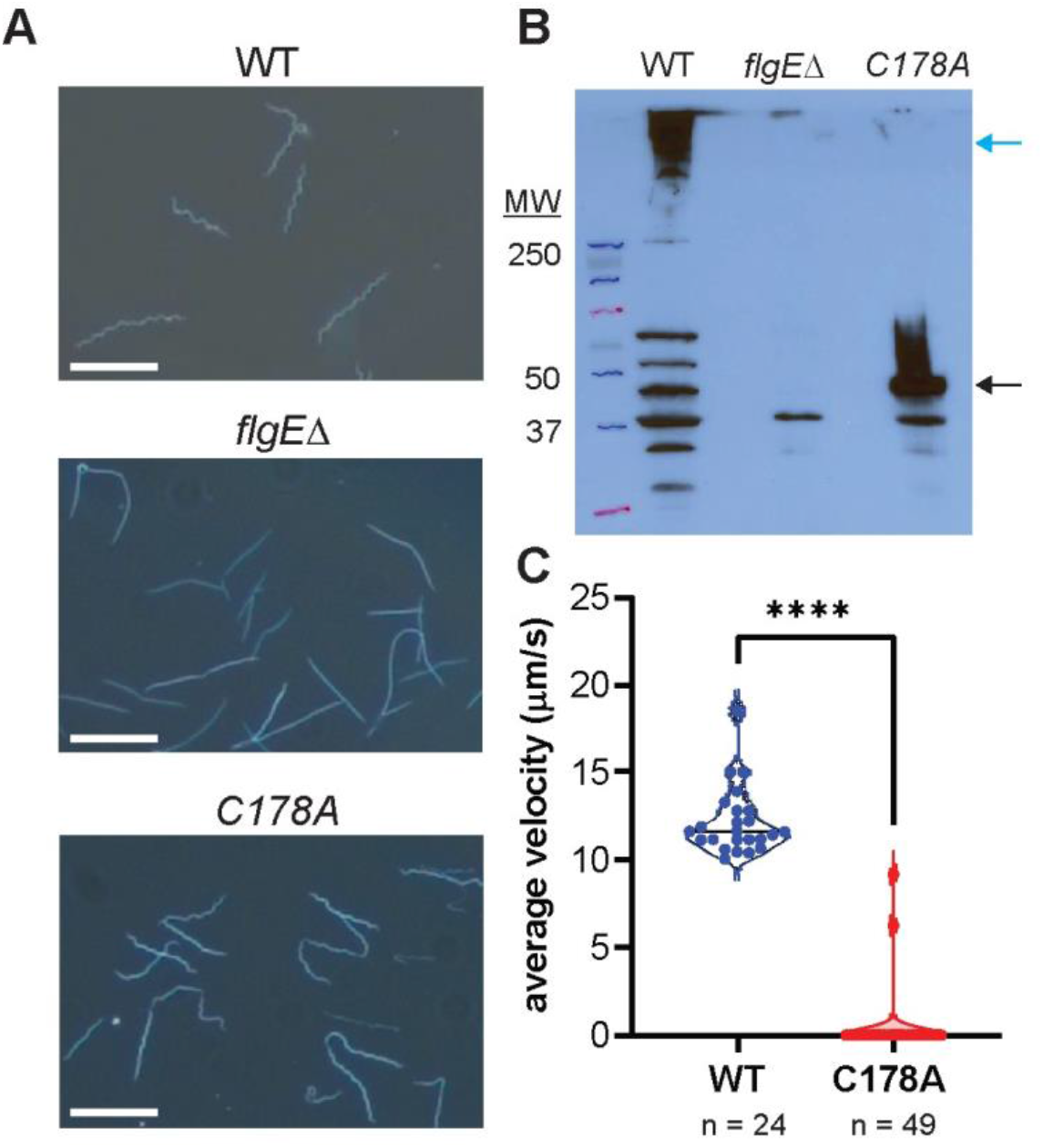
Motility of *B. burgdorferi* depends on the presence of Lal-crosslinked FlgE subunits. **(A)** Cell morphology of WT, *flgE* knock-out (*flgE*Δ), and FlgE C178A mutant cells (white scale bar = 10 μm). **(B)** Detection of FlgE HMWCs (blue arrow) and monomers (black arrow) by immunoblotting with a specific antibody against Bb FlgE. **(C)** Swimming speed analysis of individual WT (n=24) or FlgE C178A mutant (n=49) *B. burgdorferi* cells. Significance was determined via a two-tailed Student’s t-test (p< 0.0001, ****). The figure was prepared using GraphPad Prism.

## Discussion

Pathogenic spirochetes are responsible for a myriad of mammalian illnesses including Lyme disease, syphilis, periodontal disease, leptospirosis, pinta, yaws, endemic syphilis, swine dysentery, and bovine digital dermatitis. In these bacteria, motility is an essential virulence factor; loss of key flagellar genes attenuates infections for *B. burgdorferi, B. hyodysenteriae*, and *L. interrogans*.^10,24,53–55^ In addition to initial findings that FlgE crosslinking through Lal plays a critical role in the motility of *T. denticola*^36^, the Lal PTM appears to be found throughout spirochete bacteria. Furthermore, the Lyme disease pathogen *B. burgdorferi* also relies on Lal to produce functional periplasmic flagella. Using HPLC-MS coupled with electron transfer dissociation (ETD) fragmentation of the parent Lal crosslinked peptide, Lal crosslinked FlgE peptides were detected from *T. denticola, B. burgdorferi, B. hyodysenteriae, T. phagedenis, T. pallidum*, and *L. interrogans*. ETD-induced fragmentation was essential and superior to collision-induced dissociation (CID) for detecting and verifying the presence of Lal crosslinked peptides due to the tendency for Lal crosslinked peptides to assume highly charged states (e.g., +4, +5, +6, SI Figure 11). For *T. denticola* and *B. burgdoferi*, there is good agreement among *in vitro* and *in vivo*-derived FlgE samples, with Lal crosslinked tryptic peptides being confidently detected in multiple charged states in all cases. These findings allowed us to analyze Lal crosslinking in clinically relevant spirochete pathogens that would otherwise not be easily obtained (e.g., *T. pallidum*). In total, Lal crosslinking was confirmed in a selection of spirochetes belonging to three out of the four Spirochaetota orders that represent the majority of the species (Figure 1A). FlgE crosslinking was not analyzed in the smallest order Brevinematales, which is comprised of only three genera (Brevemina, Thermospira, and Longinema). Each genera contain a single species (*Brevinema andersonii, Longinema margulisiae*, and *Thermospira aquatica*, respectively), when excluding environmental, unclassified, or uncultured species.^11^ FlgE sequences are not yet available for *Longinema margulisiae* and *Thermospira aquatica*, but *B. andersonii* (Ba) FlgE maintains the conserved catalytic and/or regulatory residues required for Lal formation (Ba FlgE [accession ID: A0A1I1E638] K165, S178, and T328).

*L. interrogans* FlgE replaces the Lal precursor Cys residue with a Ser residue. Nonetheless, Li FlgE still self catalyzes the Lal crosslink. In lantibiotic biosynthesis, serine and threonine residue are dehydrated to dehydroalanine and dehydroalanine and dehydrobutyrine, respectively.^56^ However, these transformations require the activation of the Ser/Thr Oγ via transesterification or phosphorylation.^56^ The ability to form Lal from recombinant FlgE in the absence of other factors indicates that similar activation reactions are not required *per se*; nevertheless, the Li *in vitro* reaction requires high pH (10-11). Due to the replacement of Cys with Ser in Li FlgE, the reactivity of Li FlgE is markedly lower compared to Td FlgE at similar concentrations. This observation agrees with previous data that showed lower HMWC production in *in vitro* Lal crosslinking assays of a Td FlgE C178S mutant compared to WT TdFlgE.^35^ Hence, the reduced reactivity of the serine substitution in Li FlgE may be compensated by other, currently unknown cellular factors. Unlike crosslinking in all other spirochetes tested, several different Lys residues can react with Li FlgE S179 to produce Lal. This lack of specificity would seem to not solely derive from the highly basic conditions of the *in vitro* reaction (pH 11), because alternative linkages are also found in the natural sample of Li PFs, and at least one of the alternative lysine residues is highly conserved in *Leptospira*.

Why *Leptospira* would catalyze different Lal crosslinks in FlgE is currently unknown. We note that PFs from *Leptospira* spp. compared to non-*Leptospira* spp. may require enhanced flexibility that the heterogeneous crosslinking could impart. Torque measurements of *B. burgdorferi* and *T. denticola* motors yield values of ∼2700 pN·nm and ∼800 pN·nm, respectively.^57^ In contrast, *L. biflexa* motors generate a considerably higher torque of ∼4000 pN·nm^57 —^ which compare to *E. coli* motors (∼4500 pN·nm).^58^ Furthermore, unlike *B. burgdoferi* and *B. hyodysenteriae*, which contain 14-22 and 16-18 PFs, respectively, and overlap in the center of the cell body, *Leptospira* spp. contain only 2 non-overlapping PFs.^1,24^ Whereas *T. denticola* contains a comparable number of PFs (2 PFs at each cell pole) as *Leptospira*, they overlap in the cell midbody, forming a ribbon of continuous PFs from pole to pole.^30^ Motility of *Leptospira* spp.. is also more complex compared to *Borrelia* spp., *Brachyspira* spp., and *Treponema* spp..^7,24^ For non-*Leptospira* spp., asymmetric CCW/CW rotation of the PFs drives wave propagation of the cell body, thereby producing thrust to drive forward movement in the direction of the pole with motors undergoing CCW rotation.^1,7,22^ In contrast, *Leptospira* spp. form an asymmetric cell shape due to PF rotation with one end forming a left-handed helix and the other forming a hook shape.^1,7^ Thrust is produced through the combined CCW rotation of the left-handed helical cell pole and CW rotation of the cell body, moving the cell in the direction of the helical pole.^24,59^ One, if not all of these factors may favor a flagellar hook in *Leptospira* with structural properties that are different from those of other spirochetes. Alternative Lal linkages will alter the constraints at the subunit interface owing to variable spacing and coupling to tertiary structure, perhaps allowing a tuning of hook flexibility to meet the unique demands of *Leptospira* motility.

In conclusion, FlgE crosslinking via Lal is an unusual PTM that appears to be an adaptation of spirochetes associated with their unusual form of flagellar motility. The PTM is conserved across the order and in at least *T. denticola* and *B*. burgdorferi, plays an essential role in motility, a key pathogenicity determinant in these bacteria. Such a modification does not exist in humans; thus, it represents a new target for development of antibiotics against pathogenic spirochetes.

## Materials and Methods

### Sequence analysis of spirochete FlgE

Multiple sequence alignment (MSA) analysis was performed on FlgE sequences for spirochetes whose genomes have been sequenced and annotated. Sequences were retrieved using Annotree^60^ with the following search parameters: KEGG: K02390, percent identity: 30, E-value: 0.00001, percent subject alignment: 70, percent query alignment: 70. Sequences were obtained for all bacterial species whereas only spirochete sequences were analyzed. Individual spirochete FlgE sequences were filtered, and redundant sequences (both in sequence and species) were removed and aligned using Clustal Omega^61^. Weblogo^62,63^ was used to create sequence conservation graphics for *Leptospira*, non-*Leptospira*, and all spirochete FlgE.

### Cloning, expression, purification, and lysinoalanine crossinking of recombinant FlgE

For recombinant FlgE samples from *T. denticola, B. burgdorferi, L. interrogans*, and *T. pallidum*, the *flgE* gene was amplified from genomic DNA and inserted into a pet28a^+^ vector in-frame with an N-terminal His6-tag using Gibson assembly. All plasmids were confirmed by Sanger sequencing. For the expression, refolding, and purification of recombinant FlgE, the protocol described previously was followed without modification.^35^ Protein concentrations were determined using the BCA assay (Pierce, Cat. No. 23227). To form Lal crosslinked FlgE *in vitro*, 5-10 mg/mL recombinant FlgE was incubated with crosslinking buffer (50 mM Tris pH 8.5, 160 mM NaCl, 1M ammonium sulfate) for 2-5 days at 4°C. For *Leptospira* FlgE samples, crosslinking buffer was changed to 50 mM CAPS pH 11, 160 mM NaCl, 1 M ammonium sulfate.

### Spirochete cell culture

Low passage infectious clone A3-68 (wild type), a derivative strain from *B. burgdorferi* strain B31 A3, was used in this study.^21^ This strain was a kind gift from P. Rosa (Rocky Mountain Laboratories, NIAID, NIH). Cells were grown in liquid Barbour-Stoenner-Kelly II (BSK-II) medium in the presence of 3.4% carbon dioxide with appropriate antibiotic(s) for selective pressure as needed, i.e., kanamycin (300 μg/ml) for *fliK*Δ mutant.^64^ *T. denticola* ATCC 35405 strains (WT and ΔfliK) were grown anaerobically in oral bacterial growth medium containing 10% (v/v) heat-inactivated rabbit serum at 37°C as described previously.^36^ *B. hyodysenteriae* wild-type B204 serotype 2 ATCC 31212 cells were cultured in a Coy chamber using a premixed gas mixture of ∼90% nitrogen, 10% CO2 at 37-38°C in brain heart infusion broth supplemented with 10% (v/v) fetal bovine serum.^65^ *T*. phagedenis Kazan 5 cells were grown anaerobically in peptone yeast extract glucose medium supplemented with 10% (v/v) heat-inactivated rabbit serum.^66^ *Leptospira interrogans* serovar Copenhageni strain Fiocruz L1-130 cells were cultured in Ellinghausen–McCullough– Johnson–Harris (EMJH) liquid medium supplemented with 1% rabbit serum until they reached logarithmic phase at 30°C.^67^

### Periplasmic flagella (PFs) purification

Isolation of spirochete PFs was performed as described previously, with modifications.^30,54,68^ Briefly, 500 mL – 2 L of cells were grown as described above and isolated via centrifugation for 15 minutes at 8000 x g. All centrifugation steps were performed at 4°C unless stated otherwise. Cell pellets were then resuspended in cold phosphate -buffered saline (PBS) pH 7.0, centrifuged again, resuspended in ∼ 6 mL of 150 mM Tris HCl pH 6.8, 160 mM NaCl, and then centrifuged one final time. The cell pellet was then resuspended in ∼3 mL 150 mM Tris pH 6.8, 160 mM NaCl, gently rocked for 5 minutes at 4°C. Resuspended cells were then treated with 300 μL 20% (v/v) Triton X-100, which was added slowly dropwise over 10 minutes, and stirred gently for 1 hour at room temperature. Cell lysis was confirmed by darkfield microscopy, and the lysate clarified via centrifugation at 15,000 x g for 45 minutes. The crude pellet was then resuspended in 3 mL of 150 mM Tris HCl pH 6.8, 160 mM NaCl and 300 μL of 0.69 U/ μL mutanolysin in 18.2 MΩ water was added and allowed to incubate at room temperature for 1 hour and then overnight at 4°C. Samples were then centrifuged at 8,000 x g for 30 minutes, and the supernatant collected. A saturated solution of ammonium sulfate was then added dropwise to the supernatant until a final concentration of 12.5% (m/v) was obtained and stirred for 20 minutes at 4°C. To pellet the wild-type and PH PFs, the supernatant was then centrifuged at 120,000xg for 2 hours and the pellet resuspended in 1 mL of 150 mM Tris HCl pH 6.8, 160 mM NaCl. Samples were stored at 4°C for 1-2 days or flash frozen in liquid nitrogen and stored at -80°C.

### SDS-PAGE and in-gel digestion of lysinoalanine crosslinked FlgE

Crosslinked recombinant FlgE samples were directly mixed 1:1 with 2x SDS-PAGE loading dye supplemented with 100 mM DTT and heated at 90°C for 10 minutes. Denatured samples were then centrifuged briefly and electrophoresed on a 4-20% denaturing Tris-glycine SDS-PAGE gel. The gel was stained with Coomassie blue, de-stained, and washed extensively with 18.2 MΩ water. Multimer high molecular weight bands were then excised and submitted for MS. PF samples were prepared in a different manner. Briefly, PF samples were diluted 10x with acetone and incubated at -20°C for 20 minutes. Precipitated protein was then centrifuged briefly at 14,800 rpm and the supernatant was discarded. The protein pellet was then briefly dried under a stream of nitrogen for 5 minutes to remove residual acetone, washed with cold 18.2 MΩ water, and resuspended in 100 μL 8M urea. Samples were then electrophoresed according to the recombinant FlgE procedure. Excised high molecular weight bands were sliced into ∼1 mm cubes and washed consecutively with 200 μL deionized water followed by 50 mM ammonium bicarbonate, 50% (v/v) acetonitrile, and finally 100% acetonitrile. The dehydrated gel pieces were reduced with 50 μL of 10 mM DTT in 100 mM ammonium bicarbonate for 1h at 60 °C, followed by alkylation with 50 μL of 55 mM iodoacetamide in 100 mM ammonium bicarbonate at room temperature in the dark for 45 minutes. Wash steps were repeated as described above. The gel was then dried and rehydrated with 100 μl AspN at 10 ng/μL in 50 mM ammonium bicarbonate and incubated on ice for 30 minutes then at 37°C for 18 hours. Digestion was stopped by adding 30 μL 5% (v/v) formic acid. The digested peptides were extracted from the gel twice with 200 μL of 50% (v/v) acetonitrile containing 5% (v/v) formic acid and once with 200 μL of 75% (v/v) acetonitrile containing 5% (v/v) formic acid. Extractions from each sample were pooled together. The pooled sample was dried in Speedvac SC110 (Thermo Savant, Milford, MA) to 200 μL to remove the ACN then filtered with 0.22 μm spin filter (Costar Spin-X from Corning), dried to dryness in the speed vacuum and analyzed as AspN digests. Each sample was reconstituted in 0.5% (v/v) formic acid prior to LC-MS/MS analysis.

### Identification of crosslinks by nanoLC-MS/MS

The digested product was analyzed by NanoLC-MS/MS analysis at the Cornell Proteomics and Metabolomics Facility. The analysis was carried out using an Orbitrap Fusion™ Tribrid™ (Thermo-Fisher Scientific, San Jose, CA) mass spectrometer equipped with a nanospray Flex Ion Source, and coupled with a Dionex UltiMate 3000 RSLCnano system (Thermo, Sunnyvale, CA). Each sample was loaded onto a nano Viper PepMap C18 trapping column (5 μm, 100 μm × 20 mm, 100 Å, Thermo Fisher Scientific) at 20 μL/min flow rate for rapid sample loading. After 3 minutes, the valve switched to allow peptides to be separated on an Acclaim PepMap C18 nano column (2 μm, 75 μm x 25 cm, Thermo Fisher Scientific) at 35 °C in either 60-minute or 90-minute gradients of 5% to 35% buffer B (98% ACN with 0.1% formic acid) at 300 nL/min. The Orbitrap Fusion was operated in positive ion mode with nano spray voltage set at 1.85 kV and source temperature at 275°C. External calibrations for Fourier transform, ion-trap and quadrupole mass analyzers were performed prior to the analysis. Samples were analyzed using the CID and ETD toggle workflow, in which MS scan range was set to 350–1600 m/z and the resolution was set to 120,000. Precursor ions with charge states 3-7 were selected for ETD MS/MS acquisitions in ion trap analyzer and an AGC target of 3 ×10^4^. The precursor isolation width was 3 m/z and the maximum injection time was 118 ms. Precursor ions with charge states 2-3 were selected for CID MS/MS and normalized collision energy was set to 30%. All data were acquired under Xcalibur 4.4 operation software and Orbitrap Fusion Tune application v3.5 (Thermo-Fisher Scientific). All raw MS data are available upon request.

### MS data analysis

All MS and CID-ETD MS/MS raw spectra from each sample were searched using Proteome Discoverer 2.5 (Thermo-Fisher Scientific, San Jose, CA) with the Sequest HT algorithm for identification of peptides. For each spirochete species (*Treponema spp*.., *Borrelia spp*.., *Brachyspira spp*.., and *Leptospira spp*..) databases with targeted protein sequences were used for PD 2.5 database search. The search parameters were as follow: two missed cleavages for AspN or trypsin digestion with fixed carbamidomethyl modification of cysteine, variable modifications of methionine oxidation, asparagine, and glutamine deamidation, protein N-terminal Met-loss, acetylation, and Met-loss+acetylation. The peptide mass tolerance was 10 ppm, and MS/MS fragment mass tolerance was 0.6 Da. Only high confidence peptides defined by Sequest HT with a 1% false discovery rate (FDR) by Percolator were considered for the peptide identification. Confident identification of Lal crosslinked peptides was achieved by manual confirmation of MS precursor ions with high mass accuracy and their associated CID and/or ETD MS/MS spectra.

### Site-directed mutagenesis of Borreliella burgdorferi

Site-directed mutagenesis was performed using Agilent QuikChange II site-directed mutagenesis kit (Agilent, Santa Clara, CA) according to the manufacturer’s instruction. The plasmid that expresses FlgE recombinant protein was used as a template to replace Cys178 with alanine, using primers P1/P2 (see SI Table 1). All mutations were confirmed by DNA sequencing.

### Constructions of B. burgdorferi flgE deletion and site-directed mutants

The *flgE::kan* plasmid was constructed to replace the entire open reading frame of *flgE* with a kanamycin resistance cassette (*aphI*). To construct *flgE::kan*, the *flgE* upstream region, *aphI*, and the *flgE* downstream region were PCR amplified with primers P3/P4, P5/P6, and P7/P8, respectively (SI Table 1). The resultant PCR fragments were fused together with primers P3/P8. The fused PCR fragment was then cloned into the pGEM-T Easy vector (Promega, Madison, WI) generating *flgE::kan*. To delete *flgE, flgE::kan* was linearized and transformed into low passage, virulent *B. burgdorferi* strain B31 A3-68 wild-type competent cells via electroporation as previously described.^69^ The deletion was confirmed by PCR and immunoblotting analyses. The resulting mutant was designated the *flgE*Δ strain. To construct *flgE* C178A, the full-length *flgE* C178A *(flgE\**) gene and the aadA1 cassette were PCR amplified with primers P9/P10 and P11/P12 (SI Table 1), respectively. And then fused together with primers P9/P12, generating *flgE\**-aadA1. The upstream and downstream region of *flgE\** was PCR amplified with primers P3/P13 and P14/P8 and then fused together with flgE*-aadA1 by PCR using primers P3/P8. The obtained DNA fragment was cloned into the pGEM-T Easy vector. To generate the C178A replacement of *flgE*, the *flgE\**-aadA1 vector was linearized and transformed into the *flgE*Δ mutant via electroporation.^69^ The resulting complemented clones were confirmed by PCR and immunoblotting analyses. Specifically, the primer set P9/P10 was used to amplify the *flgE*\* gene and sequenced to confirm the *flgE*\* replacement instead of the wild type *flgE*. The primers for constructing these two mutants are listed in Supplemental Table 1.

### Motion tracking analysis

The velocity of bacterial cells was measured using a computer-based bacterial tracking system, as previously described.^16,70^ In brief, log-phase low passage, virulent *B. burgdorferi* strain B31 A3-68 cultures were first diluted (1:1) in BSKII medium and then 20 μl of diluted cultures were mixed with an equal volume of 2% methylcellulose with a viscosity of 4000 cp (MC4000). Then *B. burgdorferi* cells were video captured with iMovie on an Apple Mac computer. Videos are exported as QuickTime movies and imported into OpenLab (Improvision Inc., Coventry, United Kingdom) where the frames were cropped, calibrated, and saved as LIFF files. Then the software package Volocity (Improvision Inc.) was used to track individual moving cells and calculate cell velocities. For each bacterial strain, at least 25 cells were recorded for up to 30 sec.

### Structural Modeling of Leptospira FlgE

To model the orientation of FlgE monomers in *L. interrogans* hooks, AlphaFold model of Li FlgE was obtained from the AlphaFold protein structure database (uniport ID: A0A1N6RW72).^71^ Li FlgE monomers were aligned to the D1-D2 domain segments of *S. typhimurium* FlgE monomers as they are arranged in a fully assembled hook (PDB 6JZT).^72^ Structural superimposition, analysis, and images were prepared in PyMol.^73^

## Data Availability

All mass spectrometry data included in the article and/or supporting information is available upon request.

## Acknowledgments

We thank the Proteomics and Metabolomics Facility of Cornell University for providing the mass spectrometry data and NIH SIG grant 1S10 OD017992-01 support for the Orbitrap Fusion mass spectrometer. We also would like to thank Elizabeth Anderson, Qin Fu, and Ruchika Bhawal for their help with MS sample preparation, assistance in running the samples, and data analysis. This work was supported by grants from the National Institutes of Health: R01AI148844 (BRC and CL), AI078958 (C.L.), DE023080 (C.L.), R21AI163663 (E.A.W.J.), as well as Fundação de Amparo à Pesquisa do Estado de São Paulo grant 2020/02678-8 (F.J.P.) and Chemical Biology Interface NIH training grant T32GM138826 (M.D.). We would like to thank Dr. Steven J. Norris and Dr. Diane G. Edmondson for providing *in vitro* and *in* vivo-cultured Treponema *pallidum* cells for our studies. We also thank Rebekah James for technical assistance in spirochete culturing and PF purification. This manuscript is dedicated to the memory of Ronald Limberger, who first detected crosslinking of spirochete FlgE in 1994.

