## Supplemental Information for "Lysinoalanine crosslinking is a conserved post-translational modification in the spirochete flagellar hook"

### Supplemental Figures and Tables

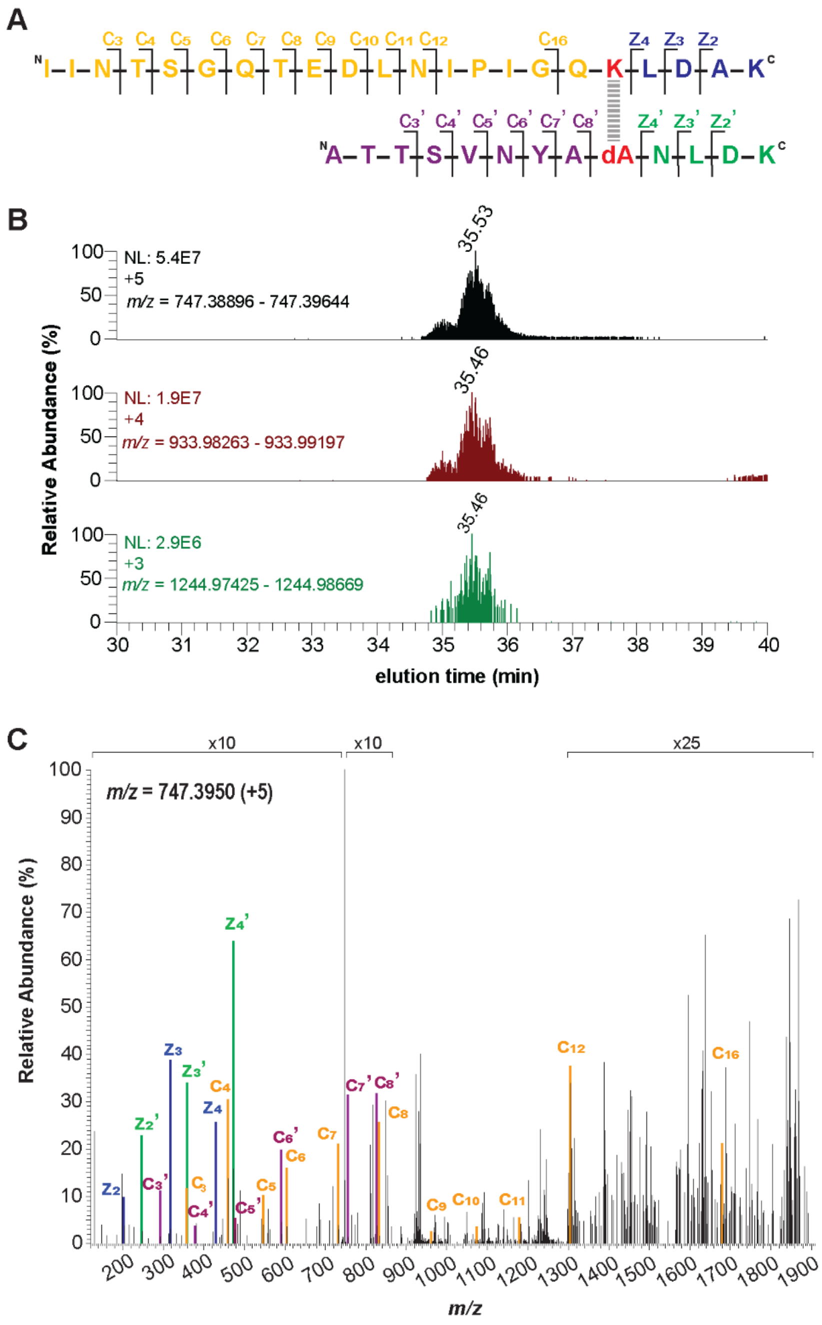


**Supplemental Figure 1:** Lal-peptide detection in *T. denticola* WT PFs. (A) Trypsin-digested Lal-containing Td FlgE peptide with c and z ions labeled as shown in (C). Lysine-165 and DHA-178 (dA) are colored red and the Lal crosslink is represented by a dotted gray line. **(B)** XICs of the +3 (top), +4 (middle), and +5 (bottom) charged Lal crosslinked peptide shown in (A). **(C)** MS/MS ETD fragmentation spectrum of Lal-crosslinked peptide +5 parent ion with c and z ions annotated and labeled according to (A). Individual c and z ions were amplified 0-25x.

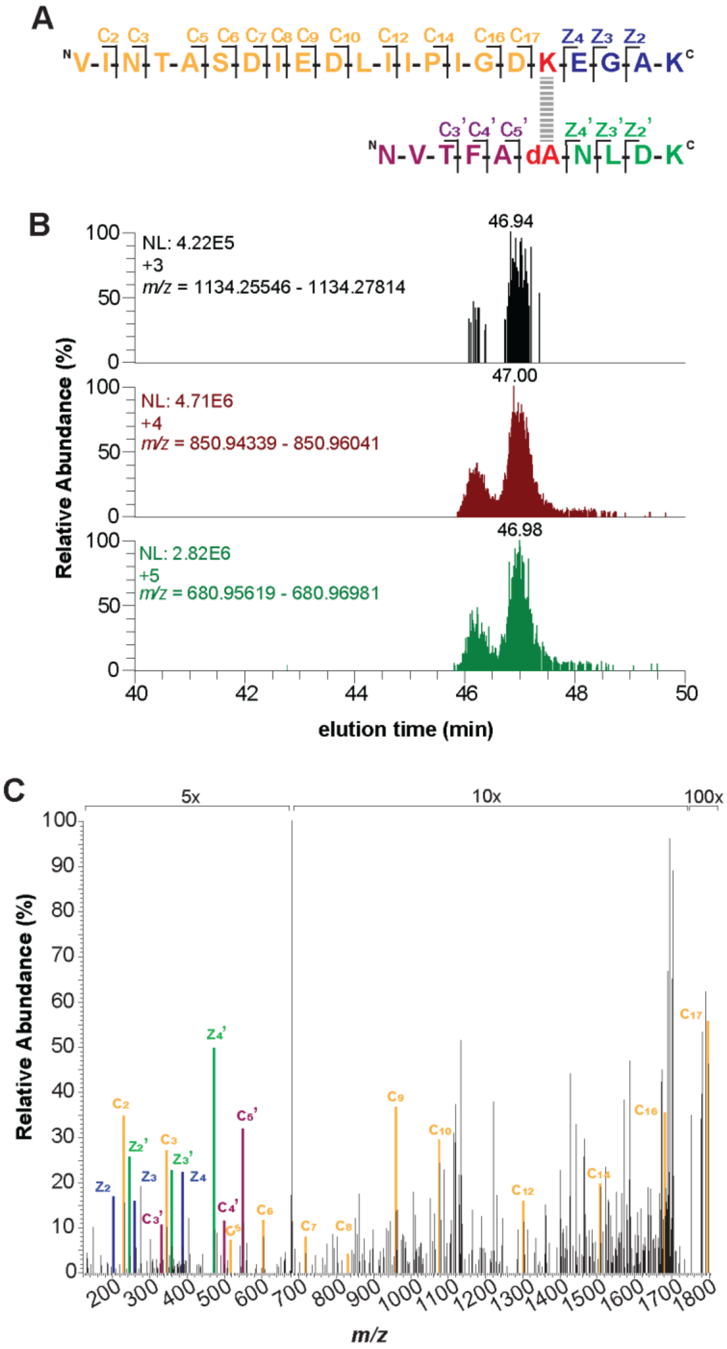


**Supplemental Figure 2:** Lal-peptide detection in *B. burgdorferi* recombinant FlgE. (A) Trypsin-digested Lal-containing Bb FlgE peptide with c and z ions labeled as shown in (C). Lysine-165 and DHA-178 (dA) are colored red and the Lal crosslink is represented by a dotted gray line. **(B)** XICs of the +3 (top), +4 (middle), and +5 (bottom) charged Lal crosslinked peptide shown in (A). **(C)** MS/MS ETD fragmentation spectrum of Lal-crosslinked peptide +5 parent ion with c and z ions annotated and labeled according to (A). Individual c and z ions were amplified 5-100x.

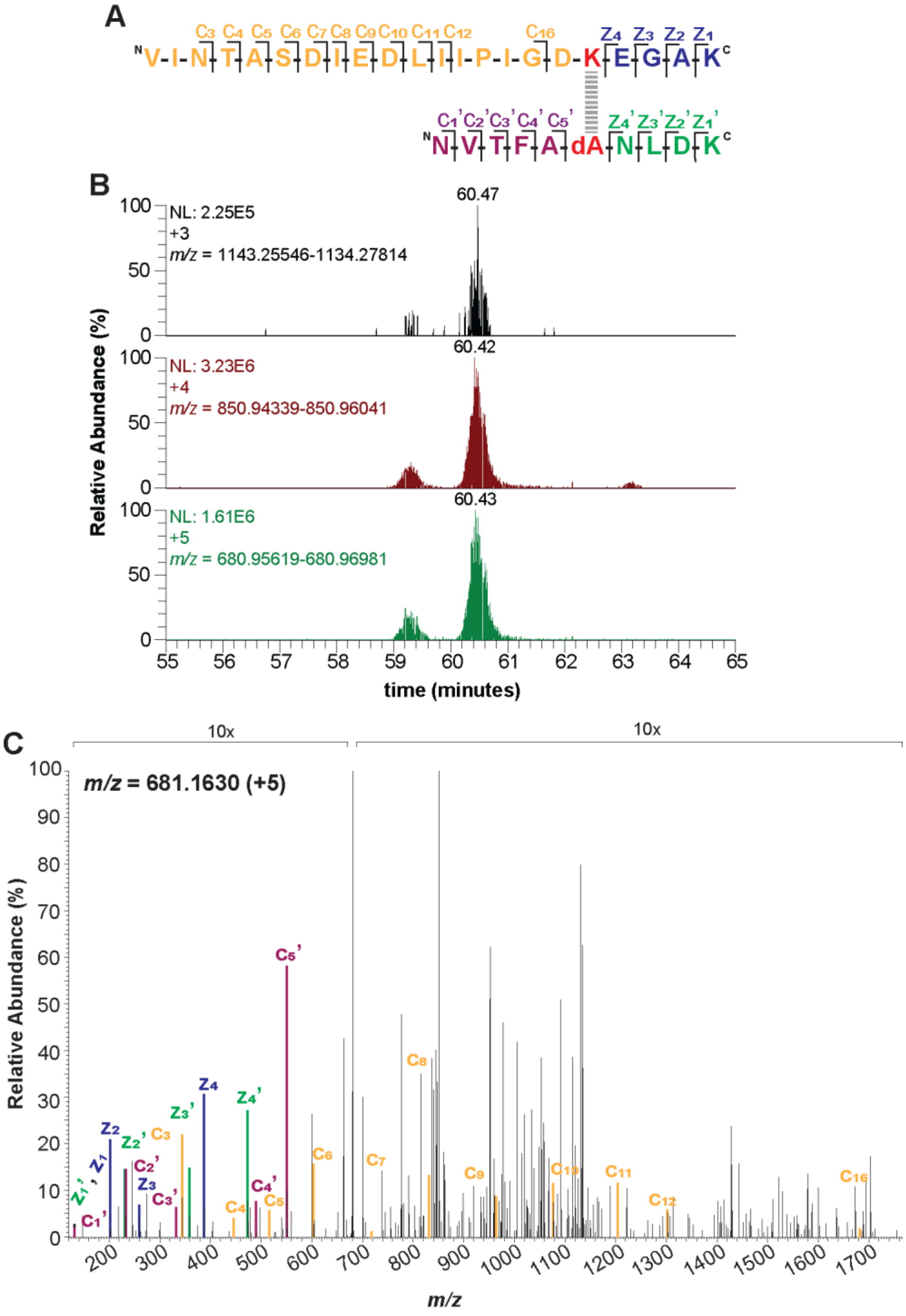


**Supplemental Figure 3:** Lal-peptide detection in *B. burgdorferi fliK*Δ PH PFs. (A) Trypsin-digested Lal-containing Bb FlgE peptide with c and z ions labeled as shown in (C). Lysine-165 and DHA-178 (dA) are colored red and the Lal crosslink is represented by a dotted gray line. **(B)** XICs of the +3 (top), +4 (middle), and +5 (bottom) charged Lal crosslinked peptide shown in (A). **(C)** MS/MS ETD fragmentation spectrum of Lal-crosslinked peptide +5 parent ion with c and z ions annotated and labeled according to (A). Individual c and z ions were amplified 0-10x.

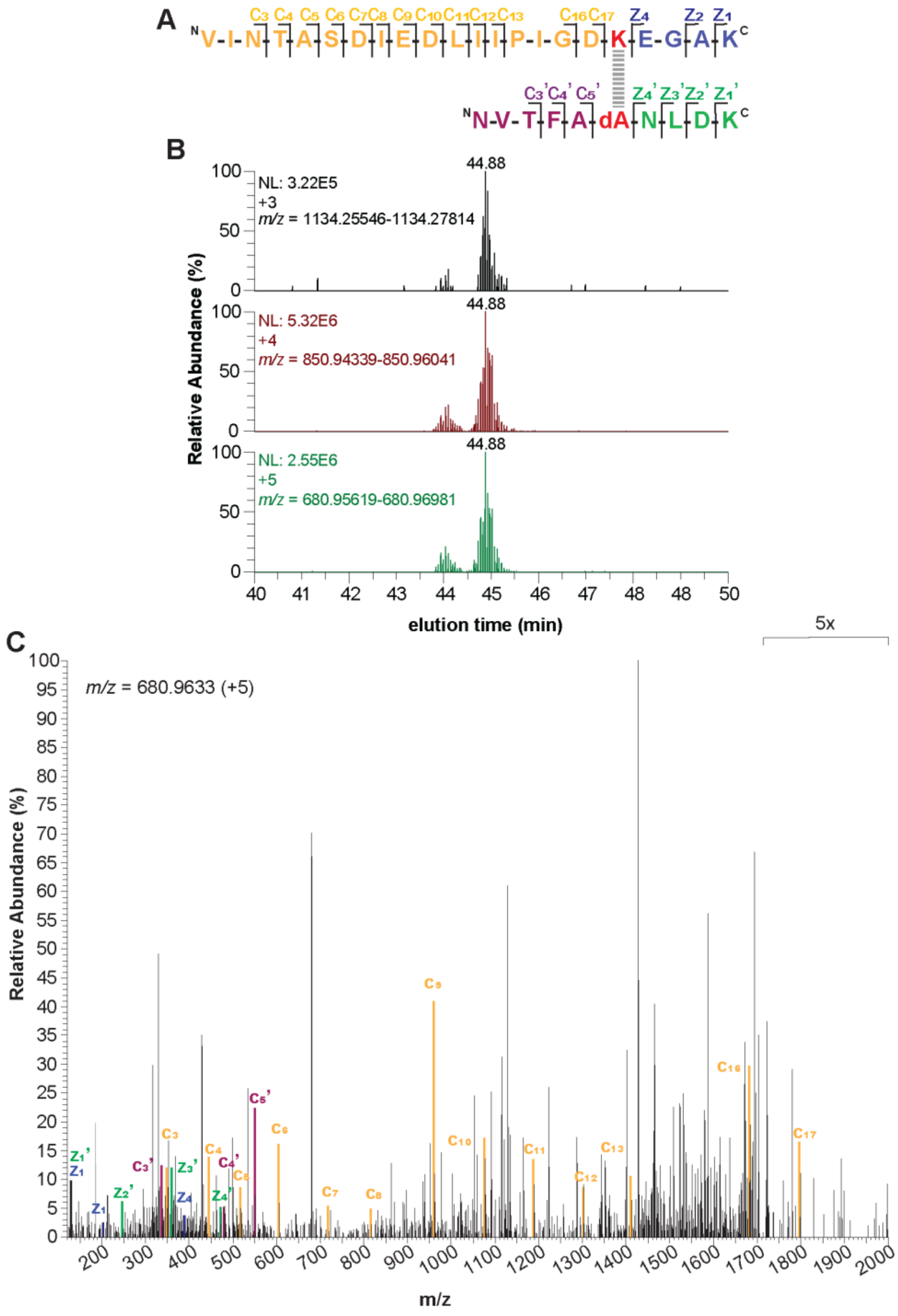


**Supplemental Figure 4:** Lal-peptide detection in *B. burgdorferi* WT PFs. (A) Trypsin-digested Lal-containing Bb FlgE peptide with c and z ions labeled as shown in (C). Lysine-165 and DHA-178 (dA) are colored red and the Lal crosslink is represented by a dotted gray line. **(B)** XICs of the +3 (top), +4 (middle), and +5 (bottom) charged Lal crosslinked peptide shown in (A). **(C)** MS/MS ETD fragmentation spectrum of Lal-crosslinked peptide +5 parent ion with c and z ions annotated and labeled according to (A). Individual c and z ions were amplified 0-5x.

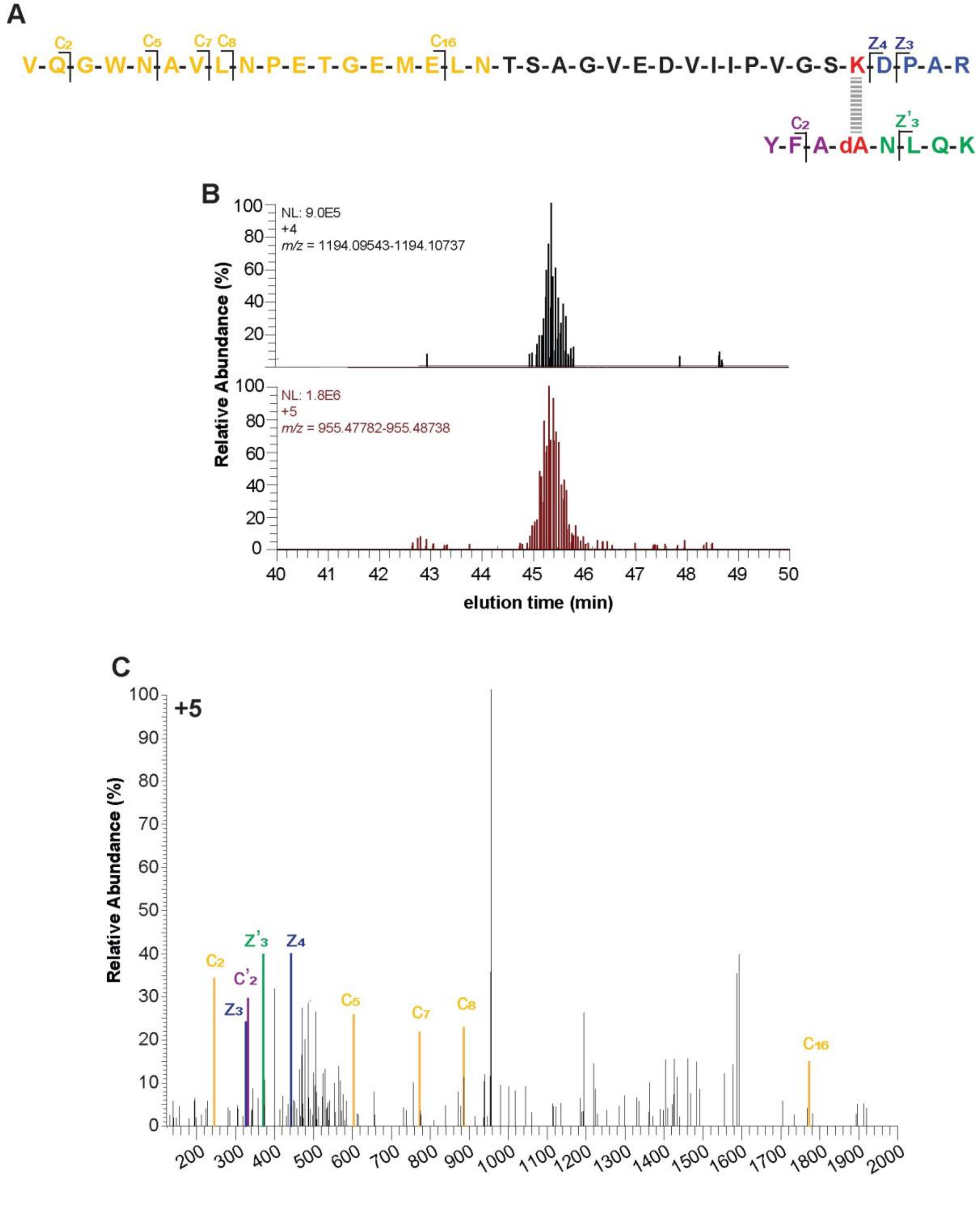


**Supplemental Figure 5:** Lal-peptide detection in *B. hyodysenteriae* WT PFs. (A) Trypsin-digested Lal-containing Bh FlgE peptide with y and b ions labeled as shown in (C). Lysine-149 and DHA-162 (dA) are colored red and the Lal crosslink is represented by a dotted gray line. **(B)** XICs of the +4 (top) and +5 (bottom) charged Lal crosslinked peptide shown in (A). **(C)** MS/MS ETD fragmentation spectrum of Lal-crosslinked peptide +4 parent ion with y and b ions annotated and labeled according to (A). Individual y and b ions were amplified 0-50x.

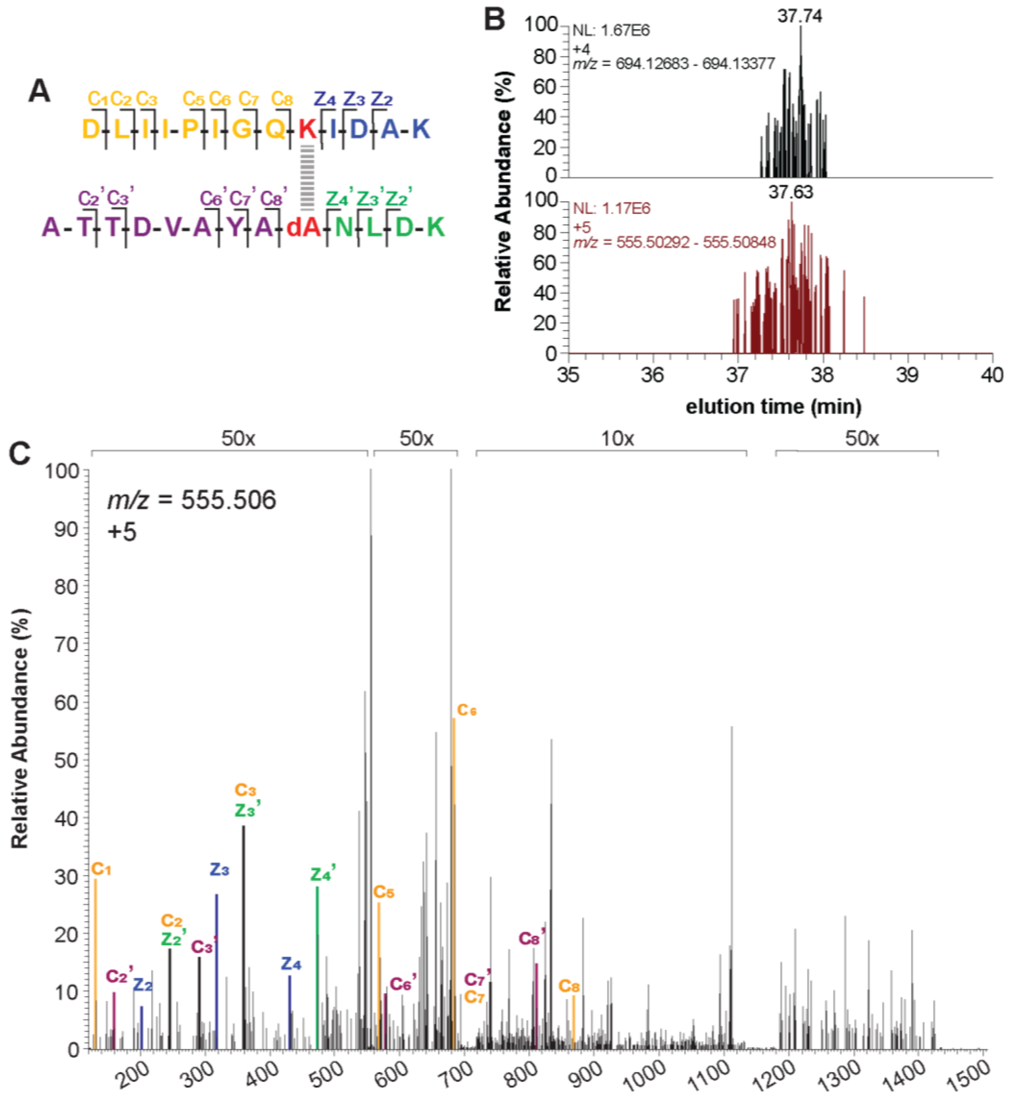


**Supplemental Figure 6:** Lal-peptide detection in *T. phagedenis flik*Δ PH PFs. (A) Trypsin-digested Lal-containing Tph FlgE peptide with c and z ions labeled as shown in (C). Lysine-165 and DHA-178 (dA) are colored red and the Lal crosslink is represented by a dotted gray line. **(B)** XICs of the +4 (top) and +5 (bottom) charged Lal crosslinked peptide shown in (A). **(C)** MS/MS ETD fragmentation spectrum of Lal-crosslinked peptide +5 parent ion with y and b ions annotated and labeled according to (A). Individual c and z ions were amplified 10-50x.

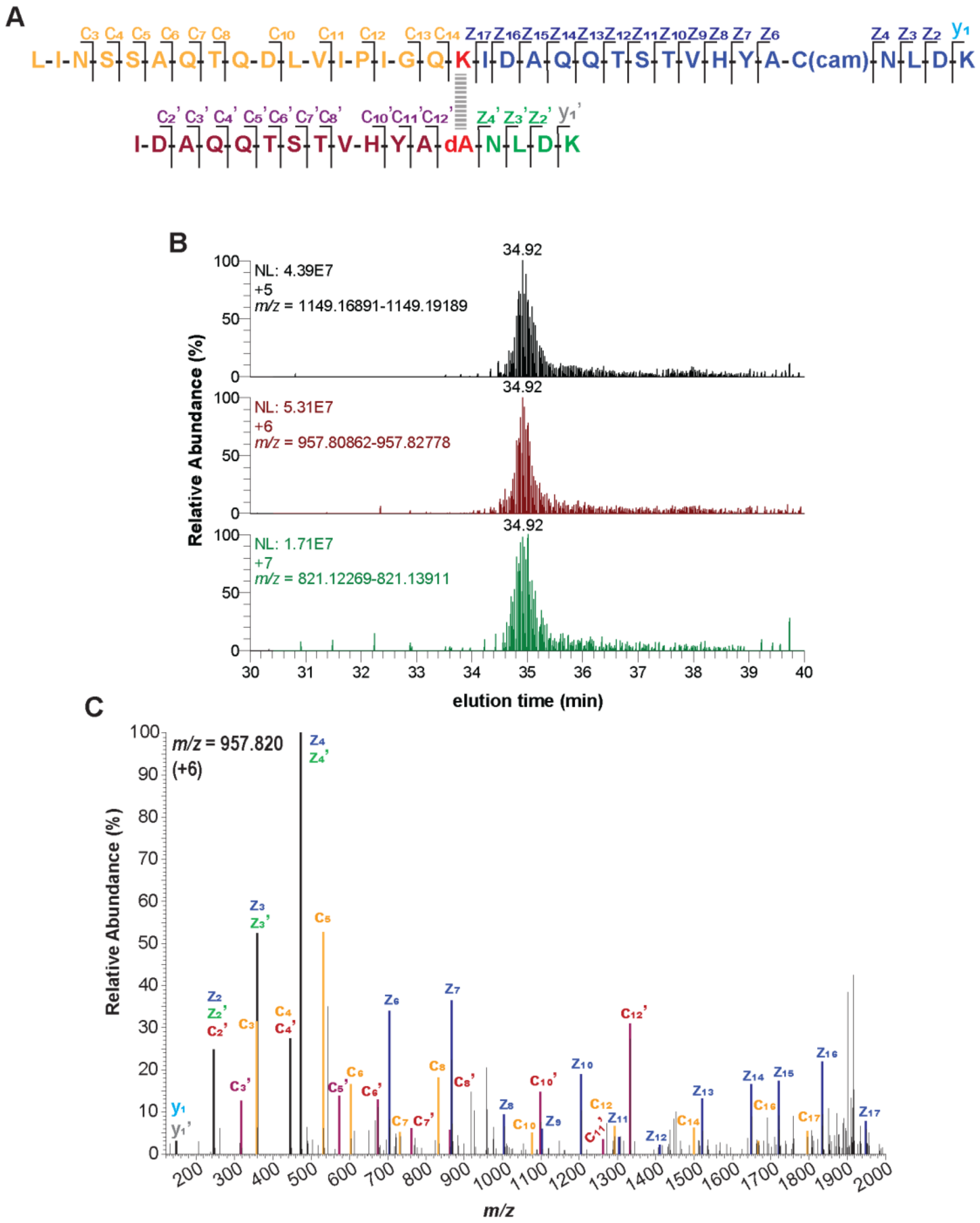


**Supplemental Figure 7:** Lal-peptide detection in *T. pallidum* recombinant FlgE. (A) Trypsin-digested Lal-containing Tpa FlgE peptide with c and z ions labeled as shown in (C). Lysine-165 and DHA-178 (dA) are colored red and the Lal crosslink is represented by a dotted gray line. **(B)** XICs of the +5 (top), +6 (middle), and +7 (bottom) charged Lal crosslinked peptide shown in (A). **(C)** MS/MS ETD fragmentation spectrum of Lal-crosslinked peptide +6 parent ion with y and b ions annotated and labeled according to (A). Individual c and z ions were amplified 0-200x.

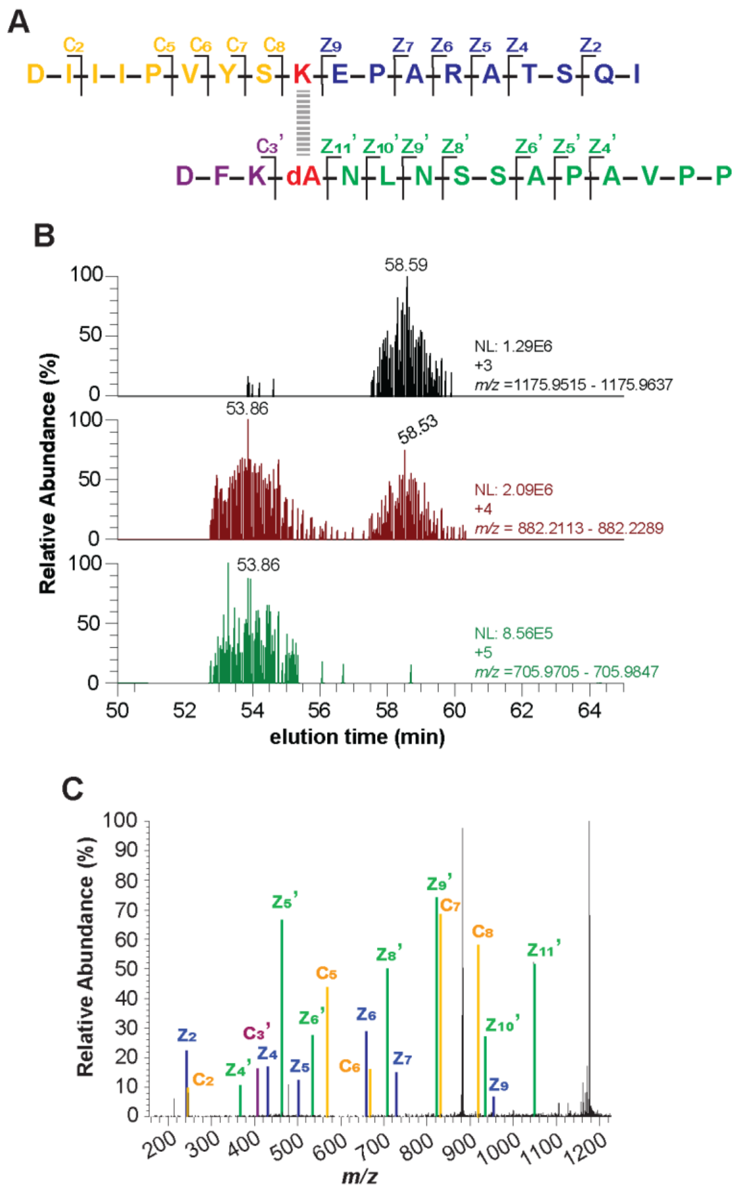


**Supplemental Figure 8:** Lal-peptide detection in *L. interrogans* recombinant FlgE. (A) AspN-digested Lal-containing Li FlgE peptide with c and z ions labeled as shown in (C). Lysine-166 and DHA-179 (dA) are colored red and the Lal crosslink is represented by a dotted gray line. **(B)** XICs of the +3 (top), +4 (middle), and +5 (bottom) charged Lal crosslinked peptide shown in (A). **(C)** MS/MS ETD fragmentation spectrum of Lal-crosslinked peptide +3 parent ion with c and z ions annotated and labeled according to (A). Individual c and z ions were amplified 0-200x.

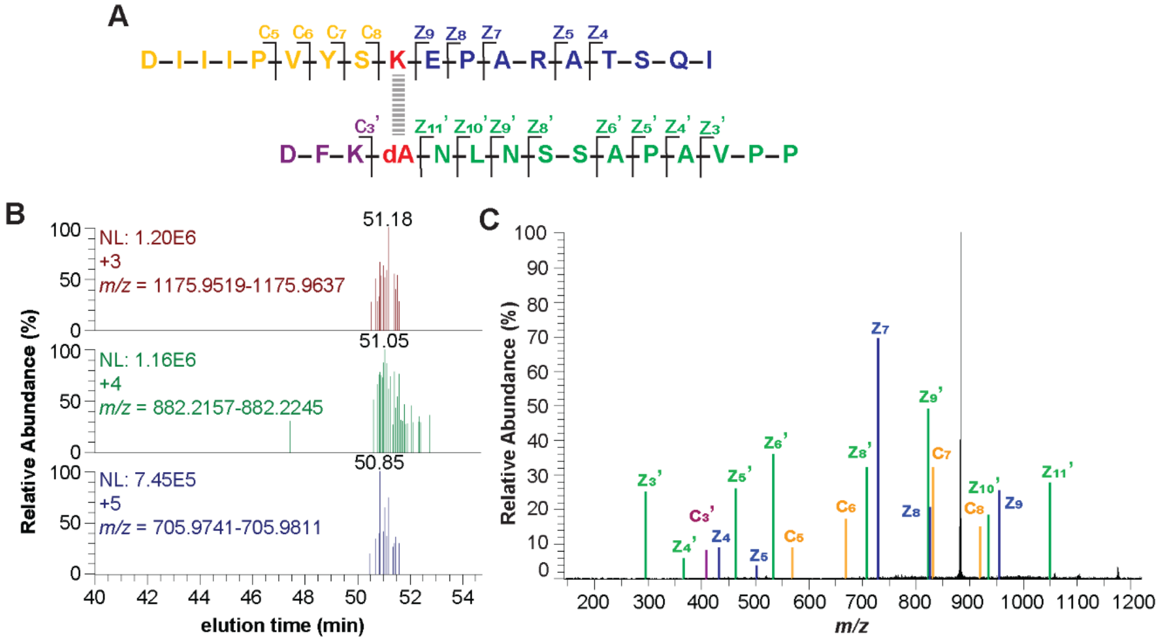


**Supplemental Figure 9:** Lal-peptide detection in *L. interrogans* WT PFs. (A) AspN-digested Lal-containing Li FlgE peptide with c and z ions labeled as shown in (C). Lysine-166 and DHA-179 (dA) are colored red and the Lal crosslink is represented by a dotted gray line. **(B)** XICs of the +3 (top), +4 (middle), and +5 (bottom) charged Lal crosslinked peptide shown in (A). **(C)** MS/MS ETD fragmentation spectrum of Lal-crosslinked peptide +3 parent ion with c and z ions annotated and labeled according to (A). Individual c and z ions were amplified 0-200x.

**A** K79/K90

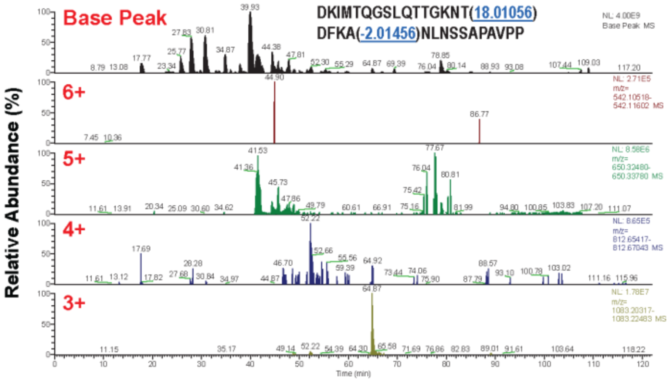


K105

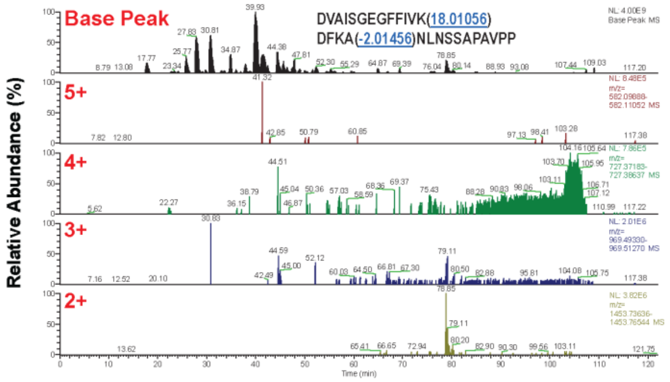


K109

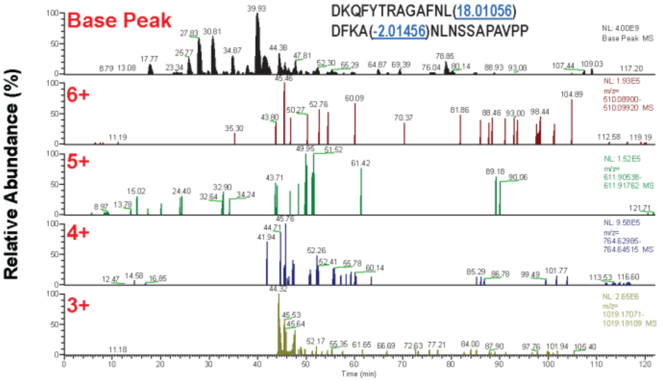


K122/K134

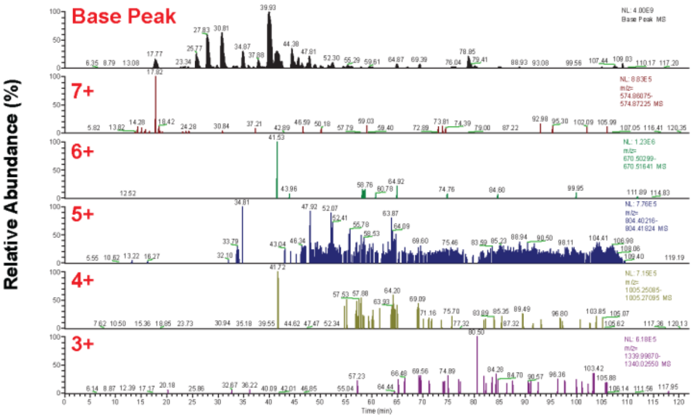

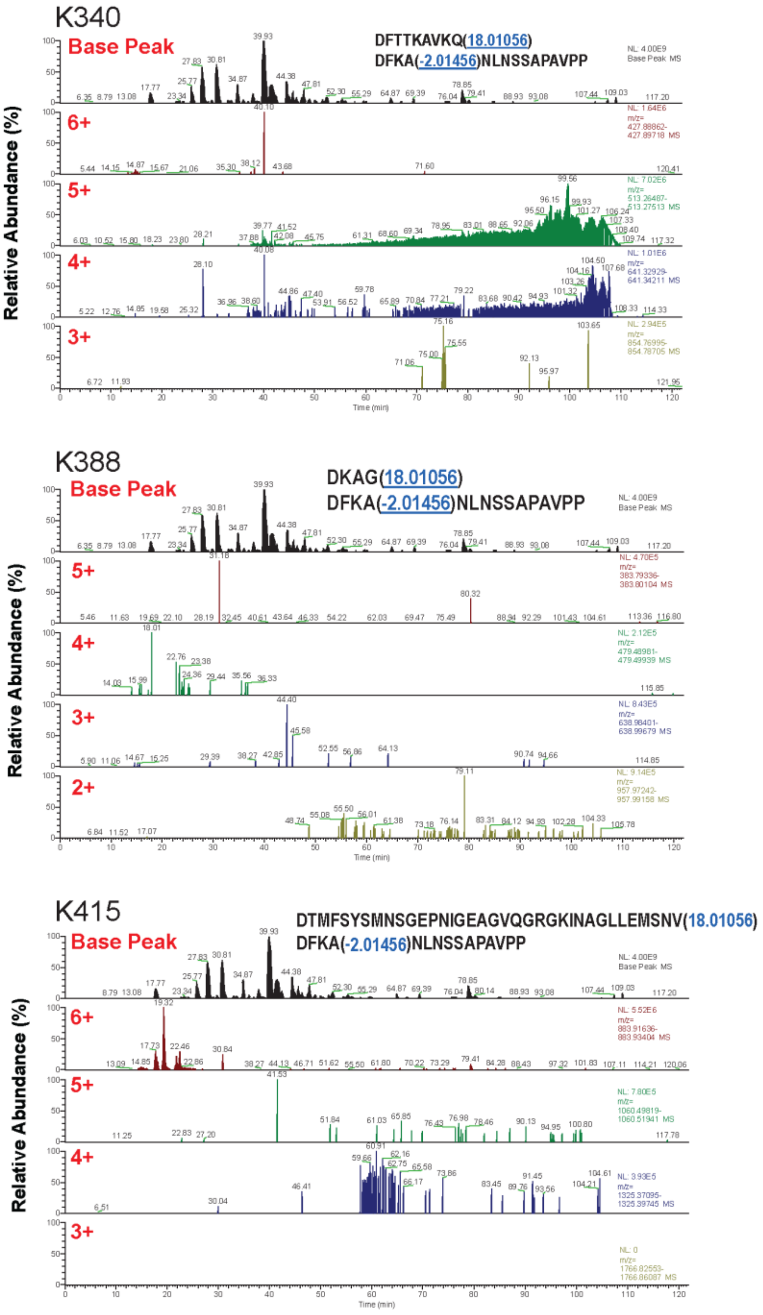
K340
Base Peak
DFTTKAVKQ(18.01056)
DFKA(-2.01456)NLNSSAPAVPP
6+
5+
4+
3+
Relative Abundance (%)
Time (min)
K388
Base Peak
DKAG(18.01056)
DFKA(-2.01456)NLNSSAPAVPP
5+
4+
3+
2+
Relative Abundance (%)
Time (min)
K415
Base Peak
DTMFSYSMNSGEPNIGEAGVQGRGKINAGLLEMSNV(18.01056)
DFKA(-2.01456)NLNSSAPAVPP
6+
5+
4+
3+
Relative Abundance (%)
Time (min)

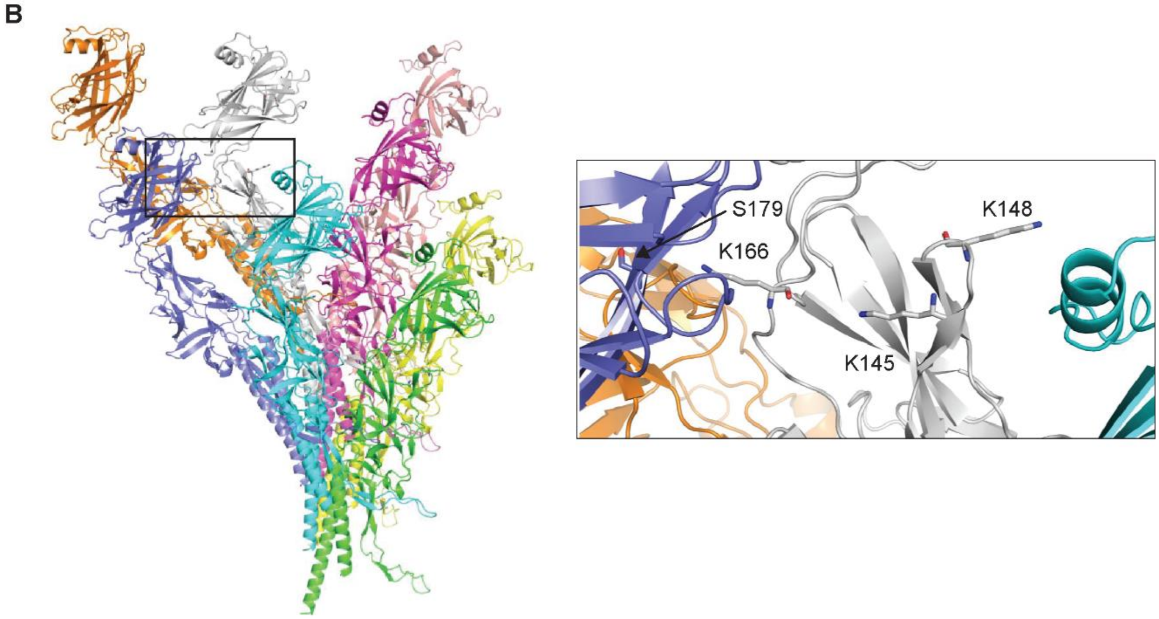


**<u>Supplemental Figure 10:</u>** <u>Analysis of other Lal crosslinks in *L. interrogans* FlgE.</u> **(A)** XICs of other potential Lal crosslinking lysine residues present in Li FlgE: Lys78/90, Lys105, Lys109, Lys122/134, Lys340, Lys388, and Lys415. **(B)** Structural modeling of LiFlgE in the *Leptospira* flagellar hook. Stick models of Lys-145, Lys-148, Lys-166, and Ser-179 residues shown for comparison.

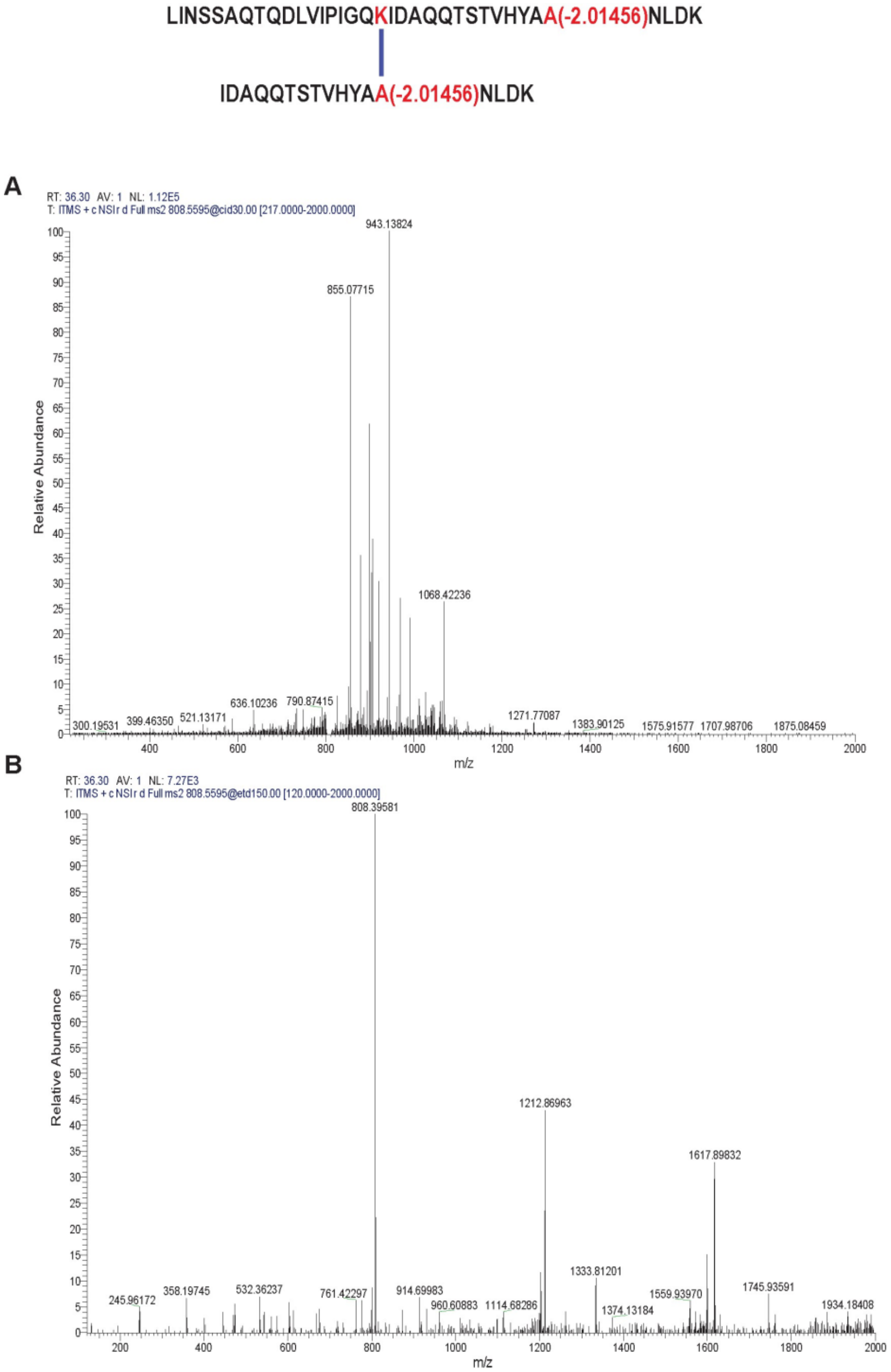


**<u>Supplemental Figure 11:</u>** <u>Comparison of CID and ETD MS/MS spectra.</u> *T. pallidum* Lal $[M+7H]^{7+}$ crosslinked peptides (see peptide above) were fragmented using either using either **(A)** CID or **(B)** ETD methods. These data reveal that ETD produced a greater number of fragment ions at higher intensity than CID for the higher charged parent Lal-crosslinked peptides.

**Supplemental Table 1.** Oligonucleotide primers used in this study

| Primers | Sequences(5'-3') | Note [a] |
|---|---|---|
| P1 | AATGTTACTTTTGCTGCTAATCTTGATAAGAGA | FlgE C178A mutation; [F] |
| P2 | TCTCTTATCAAGATTAGCAGCAAAAGTAACATT | FlgE C178A mutation; [R] |
| P3 | AAATGCTTGGTGAGAGTGGT | FlgE upstream;[F] |
| P4 | ACGTTTCCCGTTGAATATGGCTCAT ATAATTATTC CTCCAAACCT | FlgE upstream;[R] |
| P5 | ATGAGCCATATTCAACGGGA | *Kan* cassette; [F] |
| P6 | TTAGAAAAACTCATCGAGCA | *Kan* cassette; [R] |
| P7 | TTTGATGCTCGATGAGTTTTTCTAA TCTAAGATTGTTTTTTTAGT | FlgE downstream;[F] |
| P8 | TCTGCACCATCAACAACAAG | FlgE downstream;[R] |
| P9 | GT AGGTTTGGAG GAATAATTAT ATGATGAGGT CTTTATATTC TG | FlgE*; [F] |
| P10 | CTTCGGCGAT CACCGCTTCC CTCAT TTAAT TTTTCAATCT TACAAG | FlgE*; [R] |
| P11 | ATGAGGGAAG CGGTGATCGC CGAAG | aadA1 cassette; [F] |
| P12 | TTATTTGCCGACTACCTTGGTGATC | aadA1 cassette; [R] |
| P13 | ATAATTATTC CTCCAAACCT | FlgE* upstream; [R] |
| P14 | T CTAAGATTGT TTTTTTAGT | FlgE* downstream; [F] |

a * denotes the C178A; [F] forward; [R] reverse.
